# The Adhesion GPCR ADGRL2 engages Gα13 to Enable Epidermal Differentiation

**DOI:** 10.1101/2025.02.19.639154

**Authors:** Xue Yang, Feng He, Douglas F. Porter, Krassimira Garbett, Robin M. Meyers, David L. Reynolds, Duy Lan Huong Bui, Audrey Hong, Luca Ducoli, Zurab Siprashvili, Vanessa Lopez-Pajares, Smarajit Mondal, Lisa Ko, Yuqing Jing, Shiying Tao, Bharti Singal, Richard Sando, Georgios Skiniotis, Paul A. Khavari

## Abstract

Homeostasis relies on signaling networks controlled by cell membrane receptors. Although G-protein-coupled receptors (GPCRs) are the largest family of transmembrane receptors, their specific roles in the epidermis are not fully understood. Dual CRISPR-Flow and single cell Perturb-seq knockout screens of all epidermal GPCRs were thus performed, uncovering an essential requirement for adhesion GPCR ADGRL2 (latrophilin 2) in epidermal differentiation. Among potential downstream guanine nucleotide-binding G proteins, ADGRL2 selectively activated Gα13. Perturb-seq of epidermal G proteins and follow-up tissue knockouts verified that Gα13 is also required for epidermal differentiation. A cryo-electron microscopy (cryo-EM) structure in lipid nanodiscs showed that ADGRL2 engages with Gα13 at multiple interfaces, including via a novel interaction between ADGRL2 intracellular loop 3 (ICL3) and a Gα13-specific QQQ glutamine triplet sequence in its GTPase domain. In situ gene mutation of this interface sequence impaired epidermal differentiation, highlighting an essential new role for an ADGRL2-Gα13 axis in epidermal differentiation.

## Introduction

The epidermis is a self-renewing surface tissue composed primarily of specialized epithelial keratinocytes that receive signals via a variety of cell surface receptors. The balance between growth and differentiation in the epidermis represents a prototype for tissue homeostasis. Dysregulation of epidermal homeostasis contributes to numerous human clinical disorders, including cancer(1), psoriasis(2) and chronic wounds(3). GPCRs comprise the largest family of transmembrane receptors in mammals(4, 5). Several studies have addressed the role of specific GPCRs in epidermis. For example, activating mutations in the *SMO* GPCR gene are pathogenic in basal cell carcinoma(6) and the SMO inhibitor, Vismodegib, prevents cancer progression(7). Other examples include the Frizzled GPCR, which regulates keratinocyte proliferation(8), as well as LPAR GPCRs that enable epidermal differentiation(9). The roles of the many other GPCRs expressed in epidermis, however, are largely uncharacterized.

Traditional pooled CRISPR screening methods are of limited utility in characterizing the actions of gene family members in epidermal homeostasis. Standard cell survival and proliferation phenotypes within bulk cell populations can fail to detect genes with essential roles in biomedically important cell transitions, such as occurs in epithelial differentiation(10). Several CRISPR screening approaches may overcome this limitation. For example, fluorescence sorting-based pooled CRISPR screening (**<u>CRISPR-Flow</u>**), separates cells based on expression of marker proteins representative of specific cell states(11, 12), such as KRT10 for epidermal differentiation. Single cell Perturb RNA-sequencing (**<u>Perturb-seq</u>**) is another method that quantifies a subset of the transcriptome in cells undergoing a variety of genetic manipulations(13, 14). Integrating these two methods may enable systematic evaluation of the roles of members of diverse gene families in the control of homeostatic gene expression programs.

Here we performed CRISPR-Flow and Perturb-seq knockout screens of all GPCRs expressed in epidermis. These screens validated previously reported GPCR roles while also identifying a new essential requirement for the ADGRL2 adhesion GPCR (**<u>aGPCR</u>**) in epidermal differentiation. ADGRL2 expression was strongly upregulated during keratinocyte differentiation. In epidermal tissue, ADGRL2 was required for induction of a key differentiation genes involved in barrier formation. Among potential downstream G proteins, ADGRL2 selectively activated Gα13 (Gα13/Gβ/Gγ). Perturb-seq screening of genes encoding Gα subunits expressed in epidermis confirmed an essential requirement for Gα13 in epidermal differentiation. Cryo-electron microscopy (**<u>Cryo-EM</u>**) of nanodisc embedded ADGRL2-Gα13 complexes facilitated high-resolution structure determination. Structural analysis revealed an ADGRL2 ICL3 loop interface with a QQQ sequence on Gα13, which appears to be a unique GPCR-G protein interaction that may help explain the preferential engagement of ADGRL2 with Gα13. Mutation of the Gα13 QQQ sequence diminished ADGRL2 binding and impaired epidermal differentiation, underscoring the functional importance of this interface in newly uncovered pro-differentiation roles for the ADGRL2 GPCR and its Gα13 protein target.

## Results

### CRISPR-Flow and Perturb-seq of epidermal GPCRs

Analysis of dynamic epidermal differentiation RNA-seq(15) identified 101 GPCRs expressed in epidermal keratinocytes. These span the five known GPCR sub-families (**Fig. 1a**). A CRISPR library targeting these 101 GPCRs was next generated. sgRNAs were included to target known positive controls essential for epidermal homeostasis as well as non-targeting negative controls, encompassing 620 sgRNAs in total (**Extended Data Fig. S1a-b, Supplementary Table 1**). Quality control assessments confirmed minimal skew and full coverage of the resulting plasmid library (**Extended Data Fig. 1c**), which was designed to support both CRISPR-Flow and Perturb-seq screening assays. The CRISPR-Flow screen employed flow cytometry to isolate populations expressing high and low levels of the KRT10 epidermal differentiation marker for sgRNA enrichment/depletion analysis to nominate pro-differentiation and pro-progenitor GPCRs while the single-cell Perturb-seq screen evaluated GPCR gene knockout effects on keratinocyte progenitor and differentiation mRNA expression signatures (**Fig. 1b**). The complementary nature of these screens, one based on the expression of a single canonical early differentiation protein representing entry into the differentiation pathway and the other on the levels of a panel of keratinocyte mRNAs, was designed to catalog GPCRs with non-redundant impacts on homeostatic gene expression in epidermal keratinocytes.

**Fig. 1.**
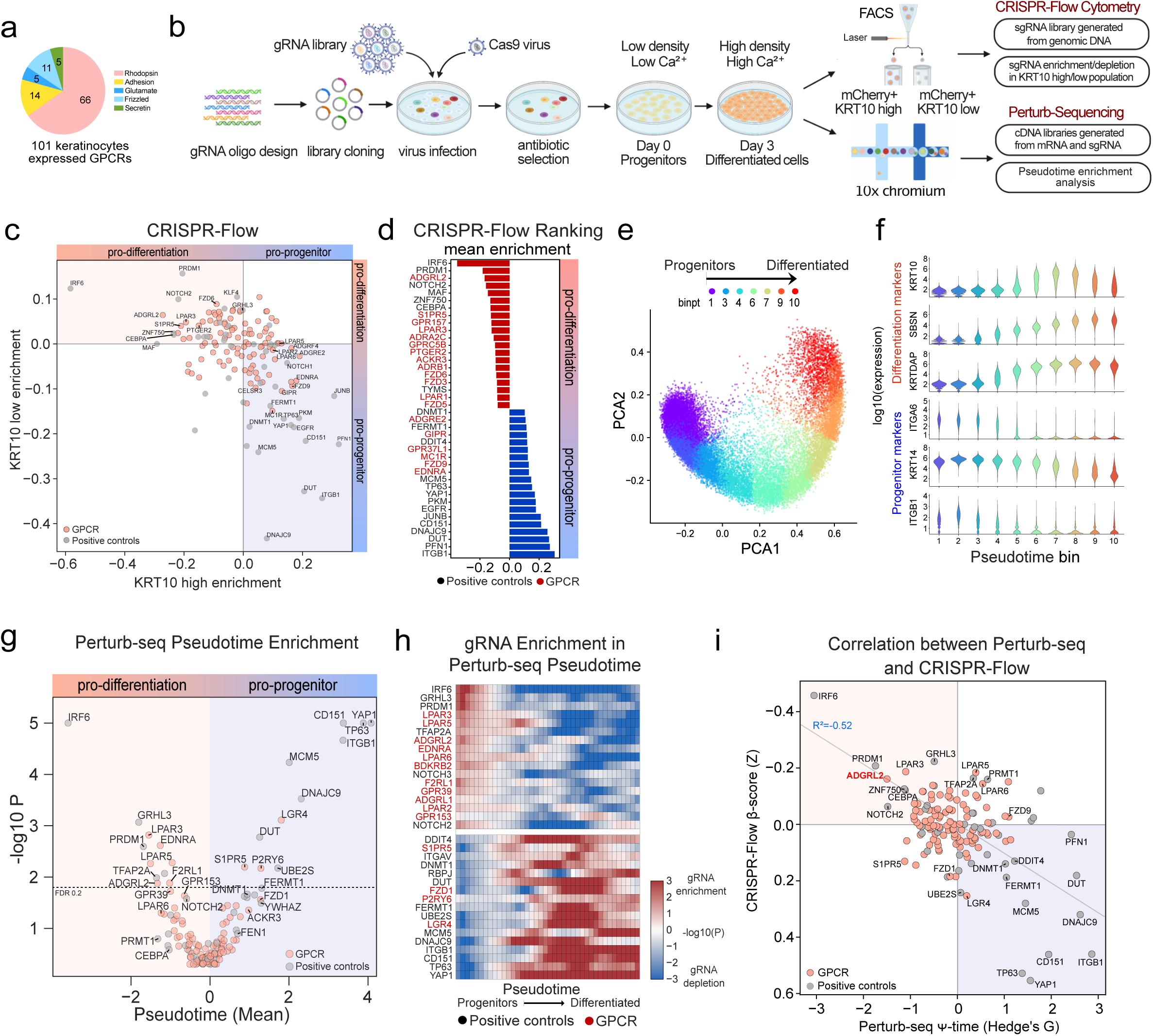
CRISPR-Flow and Perturb-seq screens in differentiating normal human keratinocytes. **a,** Pie chart of the 101 epidermal GPCRs and their classes. **b**, Schematic of experimental workflow for CRISPR screens. **c**, Scatterplot mapping guide enrichment (beta score) in low vs. high KRT10 cells from CRISPR-Flow screening. **d**, Bar graphs quantifying mean beta scores from top CRISPR-Flow identified candidates (red=GPCRs, black=controls). **e**, Principal Component Analysis (PCA) plot representing variance in gene expression between progenitor and differentiated epidermal states at day 3. **f**, Pseudotime trajectory analysis of progenitor markers ITGB1/KRT14/ITGA6 and differentiation markers KRTDAP/SBSN/KRT10 at day 3. **g**, Perturb-seq GPCR and positive control target effects plotted as pseudotime differences from non-target guides (Z score) *vs* Mann-Whitney p values for the comparison to cells with other guides. **h**, Enrichment of knock-out cells along the pseudotime trajectory (Mann-Whitney log10 P) compared to cells with safe target guides. A positive enrichment in low pseudotime cells indicates nominates a gene as necessary for differentiation. **i**, Correlation analysis of CRISPR-Flow beta scores plotted against Perturb-seq pseudotime measurements for each CRISPR target. ADGRL2 is highlighted in red. For all analyses *=p<0.05, **=p<0.01, ***=p<0.001.

Flow cytometry benchmarking for the CRISPR-Flow effort demonstrated that the KRT10-high population increased to 13.5% by day 3 of calcium-induced differentiation in vitro compared to <0.05% in day 0 progenitors (**Extended Data Fig. 1d-e**). Known positive controls required for epidermal cell differentiation, such as *IRF6, NOTCH2, PRDM1, MAF, CEBPA* and *ZNF750,* as well as the *LPAR3* GPCR, were scored by CRISPR-Flow as required for KRT10 induction. Knockout of known progenitor genes, such as *ITGB1*, *FERMT1*, *TP63*, *EGFR*, and *YAP1*, in contrast, led to increased KRT10 protein levels. These findings confirmed prior work and validated the fidelity of the screen. Among GPCRs, ADGRL2 was nominated as the strongest novel pro-differentiation GPCR (**Fig. 1c-d, Supplementary Table 2**). CRISPR-Flow screening thus validated positive controls with known impacts on homeostatic gene expression in epidermal cells while also nominating specific GPCRs as new regulators of this process.

Single-cell RNA-seq of differentiating keratinocyte populations for Perturb-seq benchmarking revealed a trajectory from progenitor-like cells to more differentiated states (**Fig. 1e**), consistent with previous studies(16). Representative differentiation genes, such as *KRT10*, *SBSN* and *KRTDAP*, as well as undifferentiated progenitor population markers, such as *ITGB1*, *ITGA6* and *KRT14*, exhibited expected expression dynamics across pseudotime (**Fig. 1f**), supporting scRNA-seq dataset quality. As an additional quality check for CRISPR-Cas9-mediated gene knockout effectiveness, expression of targeted genes was confirmed to be significantly reduced compared to non-targeted controls (**Extended Data Fig. 1f**). Perturb-seq assigned known pro-differentiation genes, *IRF6, NOTCH2/3, PRDM1, TFAP2A*, and *GRHL3*, as well as pro-progenitor genes, *ITGB1*, *FERMT1*, *TP63*, and *YAP1* to their known roles, supporting the fidelity of the screen (**Fig. 1g-h**). As in CRISPR-Flow, Perturb-seq identified specific LPAR genes as pro-differentiation GPCRs, including *LPAR3, LPAR5 and LPAR6* (**Fig. 1g-h**), which are known to be activated by Lysophosphatidic Acid (**<u>LPA</u>**) and which have been previously implicated in keratinocyte differentiation(9). ADGRL2 was again captured as a novel pro-differentiation GPCR in Perturb-seq data (**Fig. 1g-h, Extended Data Fig. 1g, Supplementary Table 3**). Correlation of the CRISPR-Flow and Perturb-seq dual screens identified ADGRL2 and LPAR3 among the top GPCR candidates promoting differentiation (**Fig. 1i**). Integration of Perturb-seq and CRISPR-Flow screens therefore nominated ADGRL2 as a novel pro-differentiation GPCR.

### ADGRL2 is essential for epidermal differentiation

In keratinocyte differentiation RNA-seq datasets(15), mRNA levels of ADGRL2 were significantly upregulated during differentiation (**Extended Data Fig. 2a, red labeled**). Consistent with this, uniform manifold approximation and projection (**<u>UMAP</u>**) analysis performed on the Perturb-seq dataset showed that ADGRL2 mRNA levels were low in progenitors but markedly elevated in differentiated cells, concordant with *KRTDAP* differentiation gene expression and inversely correlated to the *MKI67* proliferation marker (**Fig. 2a**). Further evaluation of ADGRL2 mRNA levels showed significant upregulation during differentiation by qPCR analysis (**Fig. 2b**), along with accumulation of the self-cleaved dual protein species C-terminal fragment (**<u>CTF</u>**) of ADGRL2 (**Fig. 2c**). Both ADGRL2 mRNA and protein levels were therefore significantly increased during differentiation, further suggesting a possible role for ADGRL2 in this process.

**Fig. 2.**
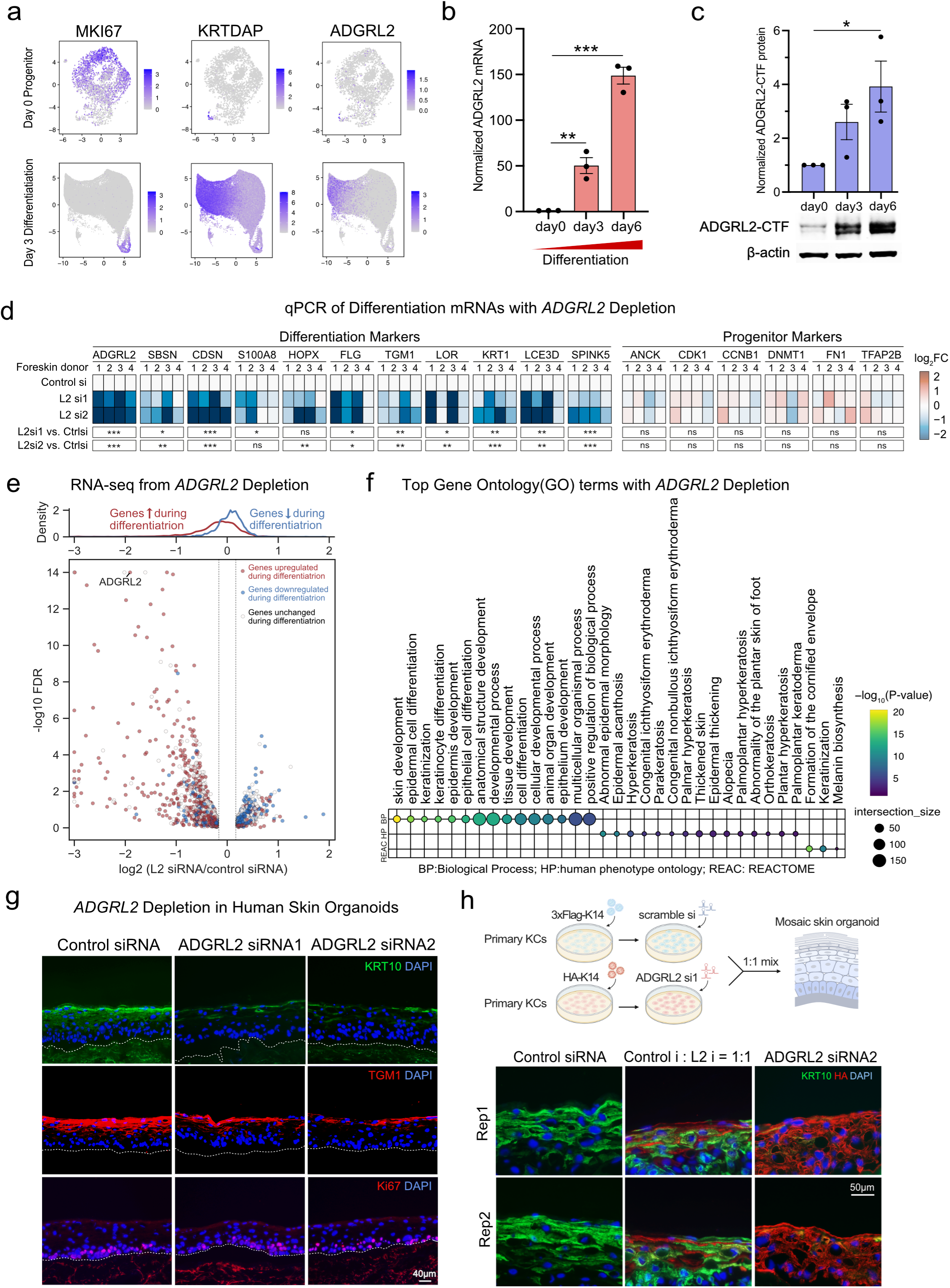
Requirement for ADGRL2 in epidermal differentiation. **a**, UMAP analysis depicting ADGRL2 expression in progenitor and differentiated states in scRNA-seq data. **b**, ADGRL2 mRNA levels across keratinocyte differentiation in vitro. **c**, ADGRL2 protein levels (C-terminal fragment dual bands) across differentiation. **d**, mRNA expression by qPCR of differentiation markers and progenitor markers with control versus ADGRL2 knockdown (L2 si1 and L2 si2) from four independent human skin donors evaluated. **e**, RNA-Seq analysis of ADGRL2 knockdown keratinocytes sourced from three unrelated skin donors, showing all genes with log2 fold change >0.2 or <-0.2, red/blue genes are the ones upregulated/downregulated during differentiation, white genes are unchanged ones. **f**, Gene Ontology analysis of all significantly altered mRNAs (FDR<0.05) following ADGRL2 knockdown based on the RNA-seq data. **g**, Immunostaining of KRT10, TGM1 and Ki67 in regenerated human skin organoid tissue with ADGRL2 depletion versus control. **h**, Immunostaining of KRT10 in mosaic skin organoids. In **b** and **c**, one-way ANOVA with Dunnett’s test against day 0 cells, n = 3. In **d**, one-way ANOVA with Dunnett’s test against control cells for each gene, n = 4.

In vitro, keratinocyte ADGRL2 depletion was associated with reduced differentiation mRNA levels as measured by both qPCR (**Fig. 2d, Extended Data Fig. 2b**) and RNA-seq (**Fig. 2e, Extended Data Fig. 2c, Supplementary Table 4**). Gene Ontology (**GO**) analysis of RNA-seq data showed that ADGRL2 depletion led to impaired induction of epidermal differentiation genes in patterns linked to human clinical phenotypes characterized by abnormalities in this process (**Fig. 2f**). Similarly, in 3-dimensional human skin organoid tissue regenerated from diploid human keratinocytes treated with siRNAs to ADGRL2 or scrambled control, ADGRL2 depletion led to decreases in differentiation proteins such as KRT10 and TGM1 in suprabasal epidermis, without affecting proliferation markers Ki67 (**Fig. 2g, Extended Data Fig. 2d**). To investigate whether ADGRL2 influences keratinocyte differentiation in a cell autonomous or non-cell autonomous fashion, mosaic epidermal tissue was generated by mixing HA-KRT14 labeled ADGRL2 knockdown cells with Flag-KRT14 labeled control knockdown cells at 1:1 ratios. ADGRL2-depleted cells displayed reduced KRT10 differentiation protein expression, in contrast to adjacent control cells (**Fig. 2h, Extended Data fig. 2e-f)**. These findings nominate ADGRL2 as a cell-autonomous pro-differentiation GPCR in epidermis.

### ADGRL2 activates Gα13

GPCRs interact with Gα subunits of heterotrimeric G proteins to induce downstream signaling. To identify specific Gα subunits activated by ADGRL2 during keratinocyte differentiation, a TRUPATH assay(17) was employed. This approach uses the BRET2 sensor to measure the attenuation of luminescence upon downstream G protein activation(17). To avoid interference by endogenous G proteins, a pan G protein protein-knockout (**<u>GKO</u>**) HEK293 cell line(18, 19) was used. Exogenously expressed ADGRL3 co-transfected with Gα13/Gβ3/Gγ9 served as a positive control and ADGRL3 co-expressed with GαoB/Gβ3/Gγ8 functioned as a negative control(20). 12 Gα proteins are expressed across keratinocyte differentiation, as detected by RNA-seq(15). A panel of 12 putative Gαβγ “epidermal transducerome” combinations was therefore evaluated in cells expressing the full-length ADGRL2, which correctly localized to the cell membrane (**Extended Data Fig. 3a**). Gα13 subunit displayed the highest basal activation levels by full-length ADGRL2 (**Fig. 3a**), even though it seems to mildly activate several other Gα proteins. The Tethered Agonist (**<u>TA</u>**) peptide serves as an intrinsic ligand in aGPCRs; once exposed by proteolytic cleavage, it activates the receptor to initiate intracellular signal transduction(21). To further dissect the role of the TA peptide in cleavage-dependent activation, therefore, two independent systems, PAR1-ADGRL2 and EK-ADGRL2, were employed. In the PAR1-ADGRL2 system, the N-terminal extracellular portion of ADGRL2 was replaced with the N-terminal domain of protease-activated receptor PAR1, creating a TA-protected variant(22). Thrombin treatment cleaves off the PAR1 N terminus, exposing a “scarred” TA peptide with a two amino acids deletion(22). Localization of the PAR1-ADGRL2 hybrid protein to the cell membrane was not abolished due to these modifications (**Extended Data Fig. 3a**). Notably, despite generating a non-native “scarred” TA peptide(20, 22), the receptor still predominantly activated Gα13 upon cleavage (**Fig. 3c**). To further study this, an EK-ADGRL2 fusion was also generated. In this approach, enterokinase-mediated cleavage selectively cleaves off the SNAP tag on the N terminus of the protein and exposed a native TA peptide(20, 23). Like the PAR1-ADGRL2 fusion, localization of the EK-ADGRL2 hybrid protein remained unchanged compared to wild-type full length ADGRL2 (**Extended Data Fig. 3a**). Upon enterokinase release of the TA peptide, the induced receptor selectively activated Gα13, reaffirming the preferential activation of Gα13 by ADGRL2 (**Fig. 3d**), and congruent with a finding reported by Pederick et al(24). Gα13 activation by these two TA peptide cleavage mimetics of ADGRL2 are consistent with the full length ADGRL2 result. However, the two engineered constructs exhibited generally weaker Gα proteins activation compared to the full-length ADGRL2 plasmid, likely because the cleavage in these constructs does not replicate the physiological native cleavage process. As a negative control, we assessed the differential dose responses of ADGRL2-CTF and its TA peptide deletion mutant (<u>Δ**7TA**</u>). This mutant has no capacity to activate G proteins(25). We also observed that the Δ7TA mutant failed to activate Gα13 in a dose-dependent fashion (**Fig. 3e**), despite similar expression and subcellular localization as ADGRL2-CTF (**Extended Data Fig. 3a-b**). Further confirmation was obtained using a Gα13 reporter SRE-RF luciferase assay, which indicated that wild-type ADGRL2-CTF displayed an enhanced ability to activate Gα13, while the Δ7TA mutant did not (**Extended Data Fig. 3c**). These data suggest that Gα13 serves as the primary Gα protein subtype activated by ADGRL2.

**Fig. 3.**
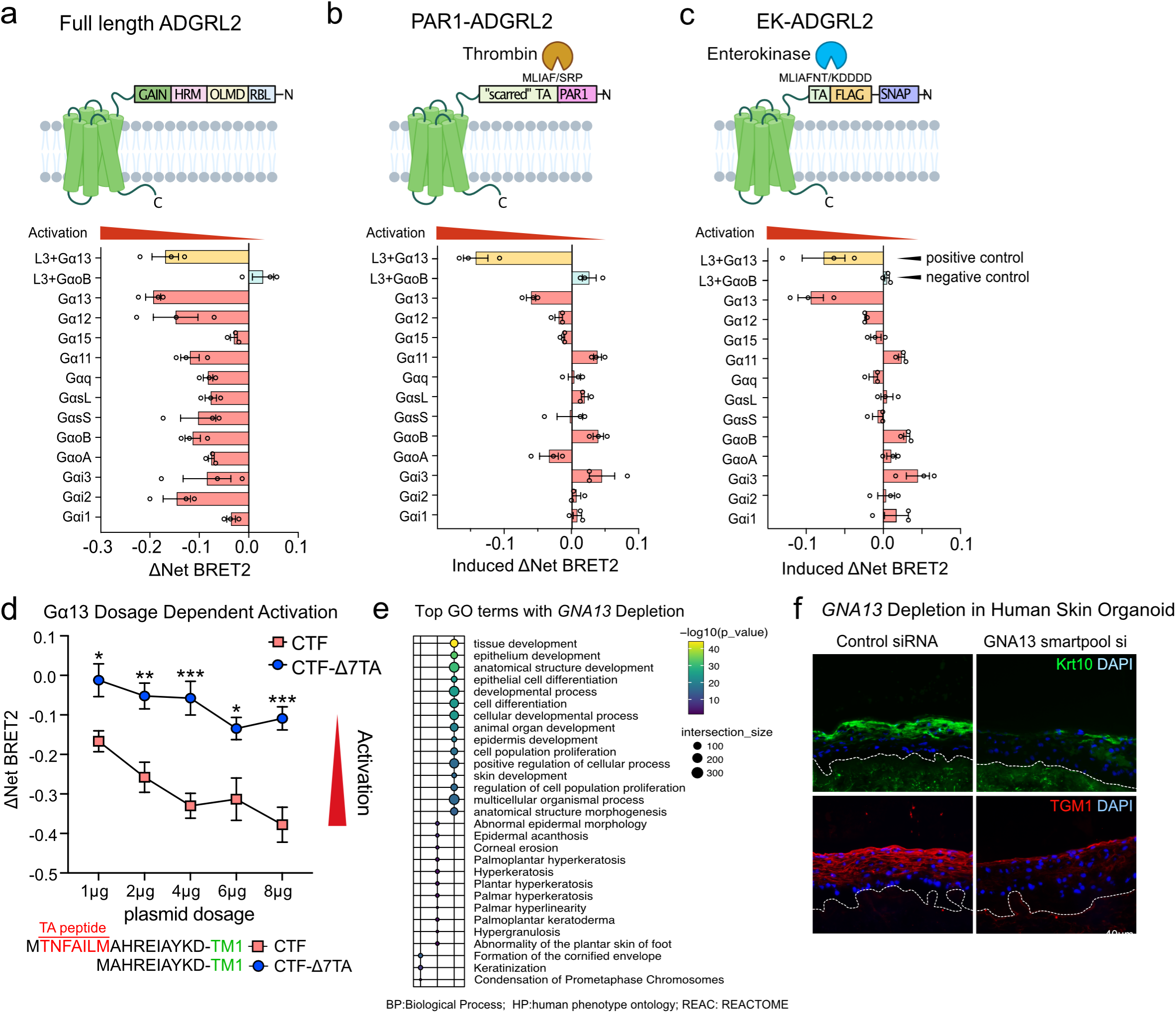
ADGRL2 activates Gα13. (**a**) Domain structures, the net BRET2 signals and summarized BRET2 heatmap from full-length ADGRL2. (**b, c**) Domain structures with drug treatment conditions, the induced BRET2 signals and summarized BRET2 heatmap from PAR1-ADGRL2 and EK-ADGRL2. **d**, Dose-dependent BRET2 signaling profiles of ADGRL2-CTF and ADGRL2 TA peptide deficient mutants (Δ7TA) on Gα13. **e**, Gene Ontology analysis of all significantly altered mRNAs (FDR<0.05) by GNA13 knockdown, based on RNA-seq data. **f**, Immunostaining for KRT10 and TGM1 in human skin organoid tissues with Gα13 depletion. In **d**, two-way ANOVA with Dunnett’s test against each dosage of wild-type ADGRL2-CTF condition, n=3.

### Gα13 is essential for epidermal tissue differentiation

To investigate if Gα13 may act in a manner consistent with a role as a downstream effector of ADGRL2 in keratinocytes, knocking down of Gα13 was performed. In 2D keratinocyte cultures, depletion of *GNA13* resulted in decreased mRNA levels of differentiation markers, as assessed by RNA-seq analysis (**Extended Data Fig. 3d-e**). GO term analysis of the RNA-seq data revealed that Gα13 depletion impaired the induction of epidermal differentiation genes, with expression patterns associated with human clinical phenotypes exhibiting defects in epidermal differentiation (**Fig. 2e**). Additionally, *GNA13* depletion was associated with a significant decrease in expression of 299 differentiation-enriched genes that are also modulated by ADGRL2 (**Extended Data Fig. 3f, Supplementary Table 5**). These findings were then further validated in regenerated human skin organoid tissue, where similar to *ADGRL2* loss, *GNA13* depletion reduced differentiation marker expression (**Fig. 3f, Extended Data Fig. 3g**). Collectively, these data indicate that Gα13, like ADGRL2, is therefore required for normal induction of epidermal differentiation genes.

### Cryo-EM structure of the ADGRL2-Gα13 complex in lipid nanodiscs

To examine the structural basis of Gα13 activation by ADGRL2, human ADGRL2-CTF and a miniGα13 heterotrimer were co-expressed in insect cells and reconstituted into lipid nanodiscs for Cryo-EM analysis (**Extended Data Fig. 4a-b**). The Cryo-EM map of the ADGRL2-Gα13 complex was determined at a global nominal resolution of 3.1 Å, with local refinements yielding further improved maps for the 7TM domain (2.9 Å) and G protein (2.9 Å) (**Extended Data Fig. 4c-f, Supplementary Table 6**). While previously reported aGPCR-G protein structures were determined in detergent micelles(21, 26–30), this ADGRL2-Gα13 complex structure provides an opportunity to examine how an aGPCR couples to G proteins in a lipid bilayer environment. Of note, two cholesterol (**<u>CLR</u>**)-like molecules were observed bound to the receptor in the Cryo-EM map, occupying hydrophobic cavities in between TM3 and TM4, and TM4 and TM5, respectively (**Extended Data Fig. 4g-h**). From a global perspective, the structure of the ADGRL2-Gα13 complex in lipid nanodiscs is not significantly different from the structure of the ADGRL3-Gα13 complex in detergent micelles(26) and shows a deeply buried TA peptide in the orthostatic 7TM binding site, bent TM6 and TM7 for G protein binding, and extensive interactions between ICL2 and the α5 helix (**Extended Data Fig. 4i-j**). Mutations of key ICL2 residues, such as the F943A and V942A/F943A mutants, dramatically reduced basal Gα13 activation by the ADGRL2-CTF without affecting protein localization and stability (**Extended Data Fig. 4k-m**), highlighting the broadly important role of ICL2 in the tethered agonist activation mechanism of aGPCRs^30^.

### ADGRL2 ICL3 in Gα13 activation

In contrast to the previously described Gα13-coupled ADGRL3 structure(26), however, the Gα13-coupled ADGRL2 structure in lipid nanodiscs revealed a distinct conformation of ICL3, which protrudes from the lipid bilayer and contacts the GTPase domain of Gα13 (**Fig. 4a, Extended Data Fig. 4n-o**). Additionally, subtle conformational differences were observed in the extracellular regions (**Extended Data Fig. 4p**). Unexpectedly, and in contrast to ADGRL3, the ICL3 of ADGRL2 appears to recognize three glutamine (**<u>QQQ</u>**) residues in the loop between helix 4 and β-strand 6 of Gα13 (**Fig. 4b**), a region we observe relatively weak EM densities suggesting the dynamic character of those elements (**Extended Data Fig. 4o**). The interactions in this region likely facilitate Gα13 coupling by ordering ICL3 and promoting the outward movement of TM6, which is positioned differently than in the ADGRL3 structure (**Fig. 4b**). Indeed, 3D variability analysis, as implemented previously(31), of the global Cryo-EM map revealed that ICL3 of the receptor and this Gα13 loop flex together (**Supplementary Movie 1**). Notably, the QQQ patch of Gα13 is unique across all Gα proteins, and is not even present in its close subfamily member, Gα12, in spite of the fact that the surrounding sequences are conserved between them (**Fig. 4c**). This uniqueness suggests the potential importance for these Gα13 residues in determining the ADGRL2 coupling specificity.

**Fig. 4.**
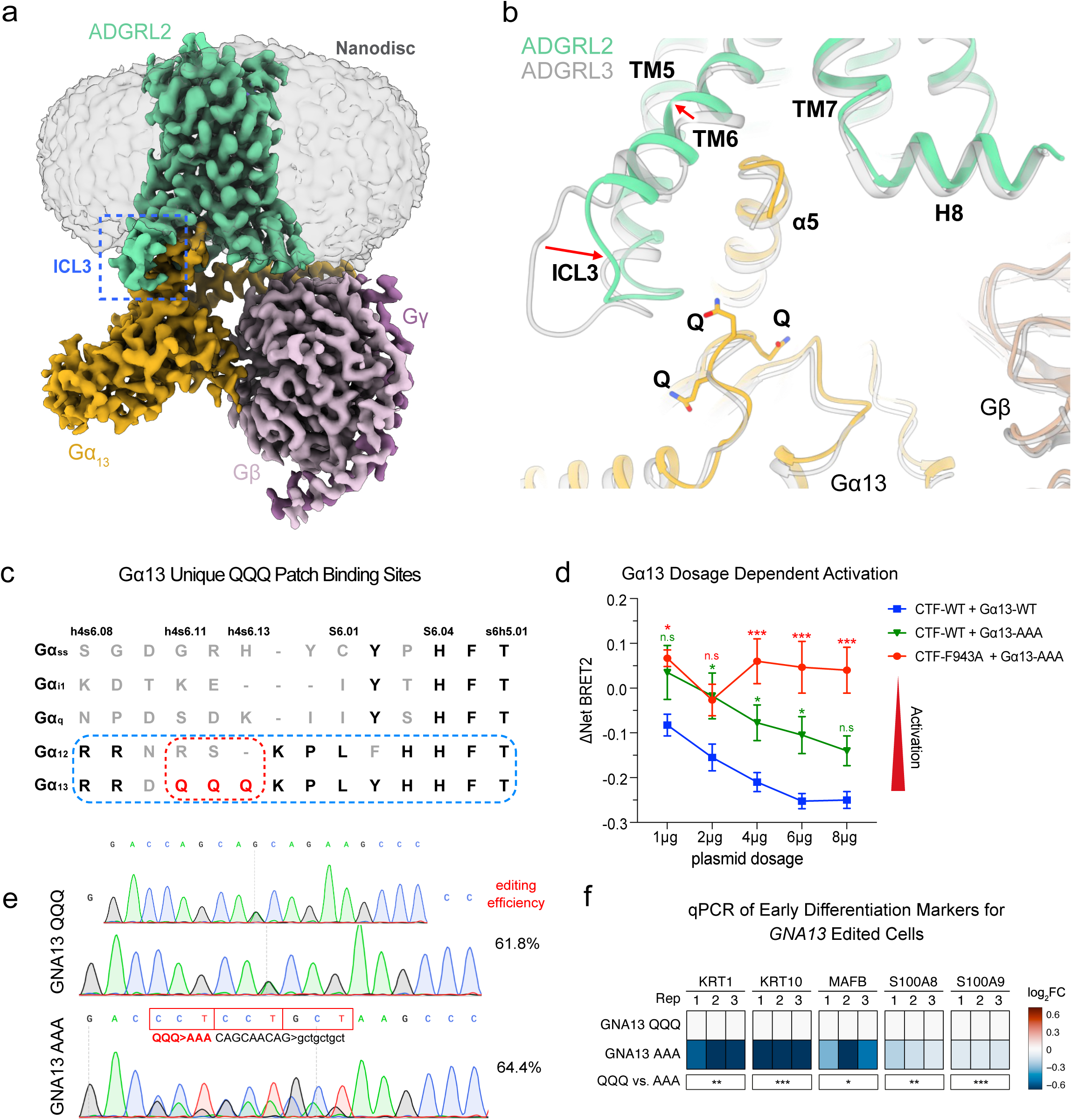
Structure of the ADGRL2-Gα13 complex. **a**, Cryo-EM map of the ADGRL2-Gα13 complex in a lipid nanodisc. The ADGRL2 ICL3 protrudes from the lipid bilayer and is highlighted with a dashed blue box. **b**, Comparison of the Gα13-coupled ADGRL2 and ADGRL3 structures showing conformational differences in ICL3 and TM6 (highlighted by arrows). In contrast to that in the ADGRL3 structure, the ICL3 of ADGRL2 is more ordered and shifts towards Gα13 to interact with a loop region containing three glutamine (QQQ) residues (shown as sticks). **c**, Sequence alignment of different Gα subtypes (including Gαs-short, Gαi, Gαq, Gα12 and Gα13) focusing on the distinct feature of three glutamine residues (QQQ) within the loop connecting helix 4 and β-strand 6 in Gα13. **d**, Effects of alanine substitution of the Gα13 QQQ triplet on the basal activity of ADGRL2-CTF WT and the ICL2 mutant (F943A). **e**, Gene editing tracks and efficiency of AAA substitution for QQQ sequences at the endogenous *GNA13* gene in bulk primary human keratinocyte populations. **f**, mRNA expression of early differentiation markers with GNA13 QQQ and AAA edited cells evaluated by qPCR. In **d**, two-way ANOVA with Dunnett’s test against each dosage of wild-type ADGRL2-CTF + wild-type Gα13 condition, n=3 or 4. In **f**, One-way ANOVA with Dunnett’s test against control Gα13 QQQ cells, n = 3.

To evaluate the importance of the novel ICL3-Gα13 interaction interface, a Gα13 mutant (QQQ to AAA) was generated and its basal activation by ADGRL2-CTF quantified. The AAA mutant exhibited similar expression levels to wild-type Gα13 but showed significantly reduced activity when activated by the wild-type ADGRL2-CTF (**Fig. 4d**). Moreover, the ICL2 mutant ADGRL2-CTF-F943A could partially activate wild-type Gα13 (**Fig. 4d**) but failed to appreciably activate the Gα13 AAA mutant (**Fig. 4d**), consistent with the premise that the ICL3-Gα13 interface contributes to Gα13 activation by ADGRL2. To examine the functional impact of disrupting the Gα13 QQQ interface with ADGRL2 on differentiation, the endogenous *GNA13* gene was edited in normal human keratinocytes to substitute AAA for QQQ residues in Gα13. Control editing was performed at the *GNA13* gene locus to alter codon sequences but preserve the QQQ residues to generate two isogenic keratinocyte populations genetically identical except for this sequence (**Fig. 4e**). AAA-edited keratinocytes displayed impaired differentiation gene induction while control-edited cells showed no such defects (**Fig. 4f**). These findings suggest that intactness of the newly identified ADGRL2 ICL3-Gα13 interface is required for normal epidermal differentiation.

## Discussion

In a prior study of skin GPCRs, RNA-seq data from keratinocyte progenitors were used to select 24 highly expressed GPCRs for individual siRNA screening, based solely on their expression levels in progenitor cells(32). However, this approach likely missed GPCRs with low expression in progenitors but significant upregulation during differentiation. For instance, ADGRL2, which exhibits minimal expression in progenitor keratinocytes, was excluded from their screening, underscoring the limitations of their screening strategy. In contrast, our study employs a more inclusive filtering method by setting a threshold of maximal transcripts per million (TPM) expression > 1 across all stages of keratinocyte differentiation and expressed in at least one squamous cell carcinoma cell line (A431, CAL27, SCC13). This broader criterion enabled us to compile a more comprehensive list of 101 skin GPCRs, providing a more robust foundation for subsequent functional screenings.

The results of the CRISPR-Flow and Perturb-seq GPCR screens demonstrated a degree of concordance, not only in identifying a novel role for ADGRL2 but also in detecting LPAR GPCRs with previously reported actions in epidermal differentiation. Some differences, however, were observed between the two screens in the candidate GPCRs identified, likely due to differences in their design. The CRISPR-Flow screen relied on differential sorting based on levels of a single canonical differentiation protein, KRT10, whereas the single-cell RNA-seq approach used in Perturb-seq assessed broader transcriptomic outcomes that were then placed within a single cell differentiation trajectory. In addition to ADGRL2, each knockout screen nominated specific GPCRs for future study as candidate regulators of epidermal homeostasis. Specific known and novel epidermal GPCRs, including S1PR5, GPR157, ADRA2C, GPRC5B, PTGER2, ACKR3, ADRB1, and FZD6, were top hits in CRISPR-Flow. Additional known and novel epidermal GPCRs, such as F2RL1, GPR39, BDKRB2, and GPR153 scored highly in Perturb-seq. Several GPCRs, such as S1PR5 and EDNRA, were assigned contrasting functions between the screens, underscoring the need for single GPCR-focused follow-up functional studies, as performed here for ADGRL2. Combined analysis of both screening methods identified more known and novel GPCRs than either screen alone, suggesting that integrating both methods may be useful in future efforts to identify new regulators of complex genetic programs.

The role of the third intracellular loop in GPCR signaling has been elusive, largely due to its high sequence variability and structural flexibility(33–35). While previous structural studies of aGPCR/G protein complexes typically relied on detergent purification methods and primarily highlighted interactions involving intracellular loop 2 (ICL2) of aGPCRs(21, 26–30), work here utilized lipid nanodiscs(36, 37), which incorporate lipid bilayers that may help recapitulate the native environment of transmembrane proteins. This may have helped to delineate the binding interface involving both ICL2 as well as ICL3 of ADGRL2 with Gα13. The present Cryo-EM structure-defined nanodisc embedded complex reveals that the ADGRL2 ICL3 engages a unique and functionally important QQQ-containing loop sequence in the GTPase domain of Gα13, an interaction which has never been observed in previous aGPCR structural studies. The QQQ site of Gα13 is unique across all Gα proteins, and is not present in its close subfamily member, Gα12, in spite of the fact that the surrounding sequences are conserved between them. Despite the noticeable conformational flexibility observed in the EM map, the ICL3 of ADGRL2 is positioned to bind Gα13’s glutamine residues, whose mutation attenuated receptor signaling and differentiation. In agreement with a recent study showing that ICL3 modulates G protein selectivity of the β2 adrenergic receptor(38), these findings suggest a regulatory role for the ADGRL2 ICL3 in tuning receptor signaling specificity. The current data represent, to our knowledge, the first reported instance of an adhesion GPCR ICL3 engaging with a Gα protein in a structured fashion, and may thus enable future potential efforts designed to modulate signaling by ADGRL2-Gα13 as well as other aGPCRs and their corresponding Gα subunit effectors.

ADGRL1, ADGRL2, and ADGRL3 reside within the latrophilin subfamily of aGPCRs(39), which have well characterized roles in synaptic function and maintenance(40, 41). While ADGRL1 and ADGRL3 are predominantly expressed in the brain(42, 43), ADGRL2 is expressed more widely(42, 43), suggesting it plays roles in other tissues. Actions of ADGRL2 in stratified epithelia, however, have been largely unexplored. Interestingly, both ADGRL2 mRNA and protein levels were significantly increased during differentiation. ADGRL2 knockdown impaired induction of a host of genes essential for epidermal differentiation, including a number whose mutation leads to human clinical disorders of this process. Among these genes are *TGM1, CDSN, LOR, SPINK5*, and *FLG*, each of whose mutation causes monogenic skin diseases characterized by abnormal epidermal barrier function, such as autosomal recessive congenital ichthyosis(44), peeling skin disease(45), keratoderma hereditaria mutilans(46), ichthyosis linearis circumflexa(47), and ichthyosis vulgaris(48) respectively. The ADGRL2 GPCR is therefore necessary for the induction of differentiation genes essential for formation of a healthy cutaneous barrier.

## Material and Methods

### Keratinocyte culture and differentiation

Keratinocytes were grown in a 1:1 ratio of Keratinocyte-SFM and Medium 154, supplemented with human recombinant epidermal growth factor (rEGF), bovine pituitary extract, and Human Keratinocyte Growth Supplement along with Penicillin-Streptomycin and Antibiotic-Antimycotic. Discarded surgically excised skin samples were incubated overnight at 4°C in a solution containing 50% dispase and 50% culture medium. The epidermis was then mechanically separated from the dermis, followed by trypsinization for 5-10 minutes at 37°C with intermittent shaking. Cellular debris and membrane remnants were removed using sterile forceps, and the cell suspension was centrifuged for 5 minutes at 1000 rpm. The resulting cell pellet was resuspended and cultured as passage 0 primary human keratinocytes. For differentiation assays, keratinocytes were seeded at densities of 1.3 x 10^6^ cells/well in 6-well plates or 300,000 cells/well in 24-well plates. The medium was fully replaced the following morning with 50:50 medium supplemented with 1.2mM CaCl_2_. Subsequent half-medium changes were carried out daily until harvesting.

### Regenerated human skin organoid tissue

Devitalized skin samples were purchased from New York Firefighter Skin Bank and subjected to an initial rinse in Phosphate-Buffered Saline (PBS). The samples were then incubated in 1x PBS containing 2x Penicillin-Streptomycin and Antibiotic-Antimycotic at 37°C for 15 days. Following incubation, the epidermal layer was separated from the underlying dermis, which was then stored at 4°C. Dermal tissue was divided into 1.5 cm square pieces and oriented within custom-designed Annular Dermal Supports (ADS) with the basement membrane matte side facing upwards and the shiny side facing downwards. Upon semi-drying, the orientation was inverted, followed by a uniform application of Matrigel to the shiny side. After Matrigel solidification, the dermal tissue was flipped. Keratinocyte Growth Media (KGM) was carefully added to the ADS plate. Keratinocytes were seeded on the basement membrane side at densities between 0.5 x 10^6^ and 1 x 10^6^ cells using a 50 μL aliquot of KGM. Media were replaced at 1–2 day intervals.

### Lentivirus generation

8 x 10^6^ Lenti-X cells were seeded in a 10 cm^2^ dish. On the following day, 5 μg of target plasmid, 5 μg of 8.91 plasmid, and 1.6 μg of pUC-MDG plasmid were transfected using Lipofectamine 3000. A complete media change was performed 6 hours or overnight post-transfection, and lentivirus was harvested 48 hours post-transfection. The viral supernatant was mixed with one-third volume of Lenti-X™ Concentrator at 1500 g for 45 minutes at 4°C. The supernatant was carefully removed, and the viral pellet was resuspended in 50 μL of cold PBS. Aliquots were prepared in small volumes and stored at −80°C for long-term use.

### Plasmid library generation

The sgRNA libraries were derived from a whole-genome sgRNA list provided by the Brunello Lab. Each gene was targeted by four unique sgRNAs. The F+E parent vector, a lentiviral expression construct, incorporates a U6 promoter followed by the sgRNA sequence. Capture Sequence 1 from 10x Genomics was integrated into the F+E plasmid downstream of the sgRNA sequence. The sgRNA sequences, flanked by the adaptors ATCTTGTGGAAAGGACGAAACACCG-sgRNAs-GTTTAAGAGCTAAGCTGGAAACAG, were synthesized by Agilent. A 200 nmol oligo library was ligated to Esp3I-digested F+E plasmid using Gibson Assembly Master Mix. Transformation was executed using Stellar Competent Cells. Transformed bacteria were plated on agar plate, and library coverage was calculated in 1:5000 dilution. All bacterial clones were harvested using 3 mL LB broth, rinsed with another 3 mL of LB broth, and then purified using the QIAGEN Plasmid Plus Maxi Kit.

### Plasmid library quality control

For the first PCR (PCR1), 50 ng of plasmid library was amplified over 5 cycles using 2x PrimeSTAR HS DNA polymerase and primers

PCR1_F:

GTGACTGGAGTTCAGACGTGTGCTCTTCCGATCGTAATACGGTTATCCACGCGG and PCR1_R: ACACGACGCTCTTCCGATCTNNNNNNNNNTGTGGAAAGGACGAAACACC to introduce a 9 bp degenerate sequence (N’s) at the beginning of read1, serving as UMIs. The PCR product was purified using the Zymo Clean and Concentrator-5 kit and further cleaned with 1.1x SPRIselect beads, followed by two 80% ethanol washes, 1-2 minutes of air-drying, and elution in elution buffer. For the second PCR (PCR2), full Illumina sequencing adapters on read1 were added using i7 and i5 primers. SYBR green was included in PCR2 and run on a Stratagene Mx3000P (Agilent) qPCR machine to monitor amplification. The PCR2 reaction was stopped before reaching plateau (around 4-5 cycles post-exponential growth). The product was cleaned using the Zymo Clean and Concentrator-5 kit and run on a 4% E-Gel™ EX Agarose Gel. The 330 bp library band was gel-purified, and its concentration was quantified using the KAPA Library Quant Kit. The plasmid library pool was run on Miseq machine (Illumina, SY-410-1003) using the MiSeq Reagent Kit v3 150-cycle kit.

### CellTiter-Blue cell viability assay

Cells were seeded at a density of 100,000 cells/well in 6-well plates. Cell Titer Blue media was prepared by combining Cell Titer Blue reagent with 5x diluted 50:50 media. Cells were incubated in 1 mL of CTB media for 45 minutes at 37°C in the dark. Fluorescence was measured at 560 nm excitation and 590 nm emission for 200 µL triplicate aliquots per sample.

### Perturb-seq single cell CRISPR screening and analysis

MOIs for Cas9 and sgRNA library lentiviruses were quantified using Lenti-X™ GoStix™ Plus. Primary cultured keratinocytes were transduced with Cas9 lentivirus and sgRNA library lentivirus (MOI ≤ 0.3). Media was replaced with 50:50 media 24 hours post-transduction, and drug selection with blasticidin (5 µg/ml) and puromycin (1 µg/ml) commenced 48 hours post-transduction, continuing for a minimum of 3 days. Subsequent to drug selection, cell viability was assessed using a Cell Titer Blue assay. Cells were seeded for differentiation 7–10 days post-transduction to optimize CRISPR cutting efficiency. Progenitor day 0 cells and differentiation day 3 cells were harvested using the 10x Genomics protocol for Single Cell Suspensions from Cultured Cell Lines for Single Cell RNA Sequencing. Cell number and viability were verified, and 40,000 cells/well were loaded onto 10x Genomics Chromium Chips for single-cell gem formation. For library generation, protocols for Chromium Single Cell 3’ Reagent Kits v3 (RevA) and Chromium Next GEM Single Cell 3’ Reagent Kits v3.1 Dual Index (RevB) were followed. Library quality and quantity were evaluated using Bioanalyzer’s high-sensitivity DNA assay. Concentrations of gene expression and feature barcode libraries were determined using the KAPA Library Quant Kit. All prepared libraries were pooled and sequenced on a NovaSeq S4 chip.

Raw 10x data was processed using Cellranger (v.3 for the larger screens, v.5 for the smaller screens) to produce h5 files, which were converted to Seurat objects. Count data was normalized with SCTransform11. Cells were filtered to only those with a single sgRNA, less than 5% mitochondrial reads and at least 200 RNAs. Dimensionality reductions were performed with the RunPCA and RunUMAP Seurat functions, except for the pseudotime analysis. Pseudotime analysis was performed by passing the SCT normalized expression data to the reduce_dimensionality function of SCORPIUS, using 10 dimensions and Pearson distances, then trajectories were inferred with SCORPIUS12. For the day 3 larger screen, the data was subset to the 2,000 most variable features (using the Seurat function FindVariableFeatures) before dimensionality reduction to reduce memory usage. To test for effects on pseudotime scores, single-sided Mann-Whitney U tests were performed on cells with guides for a target gene vs all other cells with a single guide. To analyze cell enrichment at specific points along the pseudotime axis, a Gaussian kernel density estimate (KDE) was applied to the pseudotimes of all cells, as well as to those cells targeted by guides specific to the gene of interest, using the ‘scipy.stats.gaussian_kde’ function. This KDE was then used to estimate the density of cell counts across the pseudotime axis at 40 evenly spaced points. A window of length 10 was rolled along the pseudotime space and one-sided Mann-Whitney U tests were applied to the density estimates from cells with the gene of interest vs all cells.

### CRISPR-Flow and analysis

Two biological replicates of primary human keratinocytes from separate donors were infected with lentivirus containing the CRISPR knockout library. Differentiated and progenitor cells were harvested on day 3 using trypsin for subsequent staining. Following detachment, cells were washed with PBS and fixed with 1x Fix/Perm Buffer for 20 minutes at room temperature. Cells were washed twice with 1x Perm Wash Buffer and stained with 1 µl antibody/million cells, followed by additional washes and resuspension in 1x Cell Staining Buffer. Cells were filtered through a polystyrene test tube with a cell strainer snap cap and sorted using a FACS machine. The sorted cells were then centrifuged at 800g for 5 minutes, washed with 1x PBS, and either stored at −80°C or processed immediately for genomic DNA extraction. To extract DNA, cells were resuspended in lysis buffer (50 mM Tris-HCl, pH 8.1, 10 mM EDTA, 1% SDS) and then heated to 65°C for 10 minutes with shaking to reverse formaldehyde cross-linking. RNaseA was added and samples were incubated at 37°C for 30 minutes with shaking. After that, Proteinase K (20 mg/mL) was added, and the samples were further incubated for 2 hours at 37°C, then at 95°C for 20 minutes. DNA purification was performed using SPRI beads, using 1.8x sample volume of beads. PCR amplifications were carried out using PrimeStar in 100 µL reaction volumes with 500 ng of genomic DNA and 20 pmol of each primer per reaction. The PCR program was set to an initial denaturation at 98°C for 1 minute, followed by 5 cycles of 98°C for 10 seconds, 56°C for 10 seconds, and 72°C for 15 seconds, with a final extension at 72°C. A Zymo PCR cleanup kit was used to combined PCR reactions. Subsequently, a SPRI cleanup was performed to eliminate primers from the samples. This purified product underwent a second round of PCR using PrimeStar, with the addition of 20 pmol of each primer (TruSeq adapters with dual i5/i7 indexing) and 33x SYBR Green for real-time monitoring. The PCR conditions were the same as the first, except the final 72°C elongation time for 1 minute. The reaction was stopped at cycle before plateau based on the amplification trace. PCR product was then run on a 2% agarose gel and gel purified for sequencing. Sequencing was conducted using a NovaSeq S4 chip, targeting 10 reads per cell.

Raw data was processed with a custom snakemake pipeline that counts the number of UMIs per guide in each cell pool. The number of UMIs per guide were input to MageCK RRA analysis, pairing biological samples together (“paired” option) and using the safe targeting guides as controls. Counts in the “high KRT10” group were compared against all the other divisions of that cell population (unsorted, low and medium KRT10 groups), and the reverse for the “low KRT10” group. When the p-values for the enrichment in high KRT10 vs depletion in low KRT10 (and vice versa) are combined, they are only pseudo-p-values (ψP), since each sorting group is part of the background set for the other and the P values are therefore not independent.

### siRNA reverse transfection

All ON-TARGETplus siRNAs were sourced from Horizon Discovery. A mixture of 250 pmol siRNAs and 25 µl Lipofectamine RNAiMAX was prepared in Opti-MEM and incubated for 25 minutes. Following incubation, 1×10^6^ keratinocytes were seeded in a 10 cm^2^ dish and treated with the siRNA-lipid complex for 24-48 hours.

### RNA-seq library generation and analysis

RNA was isolated using the RNeasy Plus Kit. For RNA-seq library preparation, 500 ng of RNA was processed according to the QuantSeq 3’ mRNA-seq Library Prep Kit FWD for Illumina protocol. Library quality was confirmed using Bioanalyzer’s high-sensitivity DNA assay. Library concentrations were quantified using the KAPA Library Quant Kit). Libraries were sequenced on a Novaseq S4 chip, targeting 20–30 million reads per library. Adaptors and polyA tails were removed using BBDuk (v.39.01). The processed data were aligned to the hg38 human genome (Ensembl release 99) using STAR (v.2.7.1a). Gene read counts were calculated using rsem-calculate-expression (v.1.3.1) and assembled into a data matrix via rsem-generate-data-matrix. Normalization and analysis of the RNA read count matrix were performed using the Bioconductor R package DESeq2 with default settings. The Benjamini-Hochberg method was applied for multiple testing correction to assess the false discovery rate (FDR). Gene Ontology (GO) analysis was performed using the gProfileR R package(49) according to the authors’ tutorial and identified significant GO terms associated with the most differentially expressed genes.

### Reporter assay

The SRF-RE reporter was obtained from Promega (E1350). LentiX (HEK293T) cells were plated at a density of 40,000 cells per well in 24-well plates. We transfected 100 ng of SRF-RE-Luc plasmid, 10 ng each of ADGRL2 full-length, ADGRL2-CTF, and ADGRL2-CTF-Δ7TA plasmids, along with 10 ng of pRL-Renilla plasmid (Promega) using Lipofectamine 3000. Cells were lysed 24 hours after transfection using 1x Passive Lysis Buffer (Promega). Dual-luciferase activity was measured using the Dual-Luciferase Reporter Assay System (Promega) on a Tecan Infinite M1000 instrument. Data analysis included normalizing firefly luciferase activity to renilla luciferase, expressed in Relative Luciferase Units (RLU), and adjusting for protein expression levels.

### BRET2 assay

HEK G-protein K.O cells(18) were plated on a 12-well plate at a density of 3–4x 10^5^ cells in 1 mL per well. HEK G K.O. media contained 1x DMEM plus 10% FBS, 1x Penicillin-Streptomycin with 1x MEM Non-essential Amino Acid (NeA) Solution. After 20–24 hours, cells were co-transfected with 1 µg different ADGRL2 receptors or increasing amounts ADGRL2 receptors (1, 2, 4, 6, and 8 µg) for dose-dependent BRET and TRUPATH plasmids at 1:1:1 DNA ratio (Gα13-RLuc8:Gβ3:Gγ9-GFP2) via TransIT-2020 (Mirus, MIR5400). Each condition required 97 µL of room-temperature 1x Opti-MEM. 1 µL each DNA TRUPAPH plasmid at 1 µg/µL concentration), 8 µL of 1µg/µL of the receptor plasmid/empty vector, and 3 µL of room temperature and gently vortexed TransIT-2020 reagent. The volume of the reactions was maintained consistent with an addition of empty vector. The TransIT-2020:DNA complexes mixture were gently mixed via pipetting 10 times and incubated at room temperature for 20 mins before adding drop-wise in the well. The plate was rocked gently side to side and incubated at 37oC for 24 hours before harvesting as follows: In each well, media was aspirated, and cells were washed with 1 mL warm PBS. Cells were detached with 300 µL warm Versene and incubated at 37oC for 5 mins then resuspended via pipetting 10 times. Cells were plated in a Matrigel-coated 96-well assay plate at a density of 30–50,000 cells per well in 200 µL complete DMEM containing 1x NeA. Each experimental condition was plated into three separate wells within the 96-well assay plate. BRET2 assays were performed 48 hours after transfection as follows: In each well, media was aspirated and cells were incubated in 90 µL of 1x Hanks’ balanced Salt Solution with 20 mM HEPES pH7.4 and 10 µL 100 µM Coelenterazine-400a diluted in PBS for 5 minutes. For the Thrombin treated BRET assay, followed by adding 10 μL of vehicle solution, Thrombin at 100 U/mL concentration for a final concentration of 10 U/mL, or 10 μL of Thrombin vehicle, which is 0.1% BSA in PBS. For Enterokinase treated BRET assay, followed by adding 10 μL of 5.5 U of Enterokinase in PBS or 10 μL of vehicle PBS. BRET intensities were measured via BERTHOLD TriStar2 LB 942 Multimode Reader with Deep Blue C filter (410 nm) and GFP2 filter (515 nm). The BRET ratio was obtained by calculating the ratio of GFP2 signal to Deep Blue C signal per well. The BRET2 ratio of the three wells per condition were then averaged. Net BRET2 was subsequently calculated by subtracting the BRET2 ratio of cells expressing no receptor from the BRET2 ratio of each respective experimental condition(20).

### Cryo-EM Constructs

The sequence of human ADGRL2 (isoform 2, Uniprot ID: O95490-2) comprising the TA peptide and the 7TM domain was cloned into pFastBac1 with an N-terminal haemagglutinin (HA) signal peptide followed by an additional methionine residue, and a C-terminal Strep tag. The construct of the miniGα13 heterotrimer was generated as previously described(26). A schematic description of the constructs is provided in Figure S4A.

### Protein expression

The human ADGRL2 and miniGα13 were co-expressed in Sf9 insect cells using the Bac-to-Bac Baculovirus Expression system (Invitrogen). Cells were grown in serum-free ESF 921 medium (Expression Systems) to a density of 3.5-4 million cells per ml and then co-infected with 1% culture volume of P2 baculoviruses for ADGRL2 and miniGα13. Cells were collected by centrifugation 48 h post infection and the pellets were snap-frozen in liquid nitrogen and stored at −80°C until use.

### Purification and reconstitution of the ADGRL2-Gα13 complex into lipid nanodiscs

Cell pellets were thawed and resuspended by Dounce homogenization in buffer containing 20 mM HEPES pH 7.5, 100 mM NaCl, 10 mM MgCl_2_, 10% glycerol, protease inhibitors and 0.1 mM TCEP. Following the addition of 25 mU/ml apyrase, the cell suspension was incubated at room temperature for 2 h. Membranes were solubilized in 1% n-dodecyl-β-maltoside (DDM, Anatrace), 0.1% cholesteryl hemisuccinate (CHS, Anatrace), and incubated for 2 h at 4°C. The supernatant was isolated by centrifugation at 100,000g for 40 min, and the solubilized proteins was incubated with Ni-NTA affinity resin in the presence of 20 mM imidazole for 1 h at 4°C with gentle stirring. The resin was packed into a gravity column and washed with 30 column volumes of buffer containing 20 mM HEPES pH 7.5, 100 mM NaCl, 10% glycerol, 5 mM MgCl_2_, 0.02% DDM, 0.002% CHS and 20 mM imidazole. The complex was eluted in buffer containing 250 mM imidazole, and then incubated with Strep-Tactin^®^XT 4Flow^®^ resin (IBA) for overnight at 4 °C with gentle stirring. The resin was collected and washed with buffer containing 20 mM HEPES pH 7.5, 100 mM NaCl, 10 mM MgCl_2_, 5% glycerol, 0.02% DDM and 0.002% CHS. Protein was eluted from the column using Strep-Tactin^®^XT elution buffer (IBA) plus 0.02% DDM and 0.002% CHS. Eluted protein was concentrated and injected onto a Superose 6 column with buffer containing 20 mM HEPES pH 7.5, 100 mM NaCl, 2 mM MgCl_2_, 0.02% DDM and 0.002% CHS. The peak fractions were pooled and concentrated. The lipid mixture (DOPC: DOPG at a molar ratio of 3:2) used for reconstituting the complex into lipid nanodiscs was prepared as previously described(37). The purified complex was mixed with the lipid mixture and the membrane scaffold protein (MSP1D1, Sigma) at molar ratio 1:150:1.5 and incubated for 1 h on ice. Bio-Beads SM2 (Bio-Rad) were then added into the mixture (0.35 g of beads per ml) and incubated with gentle rocking for overnight at 4°C. The reconstitution mixture was spun down to remove the Bio-Beads, and the supernatant was applied onto a Superose 6 column with buffer containing 20 mM HEPES pH 7.5, 100 mM NaCl, 1 mM MgCl_2_. The peak fractions containing the complex were collected and concentrated for preparing Cryo-EM grids with parallel examination by negative stain EM(50).

### Cryo-EM data collection and processing

The ADGRL2-Gα13 complex (3 μl at 5.5 mg ml^−1^) was applied to freshly glow-discharged 300-mesh R1.2/R1.3 UltrAuFoil holey gold grids (Quantifoil) under 100% humidity at 4°C. Excess sample was blotted away, and the grids were subsequently plunge-frozen into liquid ethane using a Vitrobot Mark IV (Thermo Fisher Scientific) and stored in liquid nitrogen. Frozen grids were imaged at cryogenic temperatures with a Titan Krios G2 (Thermo Fisher Scientific) Transmission Electron Microscope with a Selectris X post-column energy filter operated at 300 kV on a Falcon 4i direct electron camera at a pixel size of 0.75 Å. Micrographs were recorded with defocus values ranging from −0.5 μm to −1.5 μm using EPU (Thermo Fisher Scientific). Micrographs were recorded for 4 s with a total exposure dose of 50 electrons·Å^−2^.

For detailed breakdown of data processing and exact particle numbers, see Fig. S4C-F. Briefly, dose-fractionated image stacks were imported into RELION(51) and subjected to motion correction with MotionCor2(52). Contrast transfer function parameter estimation was performed with CTFFIND-4.1(53). Particle selection and extraction was performed with the Laplacian autopicker function in RELION. The extracted particles were imported into CryoSPARC(54) and subjected to multiple rounds of reference-free 2D classification, followed by iterative rounds of 3D ab initio reconstruction with multiple classes and 3D heterogeneous refinement to remove particles from poorly defined 3D classes. Selected particles were re-imported to RELION and then subjected to 3D refinement in RELION followed by Bayesian polishing. Polished particles were then imported back to CryoSPARC for nonuniform refinement, global CTF refinement, and local refinement with a soft mask around 7TM or miniGα13.

### Model building and refinement

For model building and refinement, composite maps were generated in Chimera(55) by merging local refinement maps of 7TM and miniGα13 using the ‘vop maximum’ command. The ADGRL3– Gα13 structure (PDB ID: 7SF7) was used as the initial model for docking into the Cryo-EM maps in Chimera. The model was then subjected to iterative rounds of manual adjustment in Coot(56) and real-space refinement in Phenix(57). The model statistics were validated in MolProbity(58). The refinement statistics are provided in Table S1. ChimeraX(59) was used for figure preparation.

### Western blot

Proteins were lysed in RAPI buffer with protease inhibitor and quantified using the BCA assay. 10-20μg of cell lysates were loaded per lane on an SDS-PAGE gel and transferred to a nitrocellulose membrane at 4°C. The resulting membrane was blocked with LI-COR blocking buffer (TBS) at room temperature for 1 hour. The membrane was then incubated with primary antibody at 4°C overnight. Membranes were washed with TBST and incubated with secondary goat anti-mouse and goat anti-rabbit antibodies (LI-COR Biosciences) at a dilution of 1:15,000 for 1 hour at room temperature. After that membranes were washed with TBST and visualized and quantified using the Odyssey CLx Infrared Imaging System (LI-COR Biosciences).

### Immunostaining

7 μm tissue sections were prepared with Cryostat HM525 NX. The organoids tissues were fixed in cold methanol. Cultured cell were fixed with 4% paraformaldehyde. The samples were incubated with primary antibodies at 4°C overnight, and following by the secondary antibodies room temperature for 1 hour. Samples were mounted in Duolink In Situ Mounting Medium with DAPI. All images were taken and processed using Zeiss fluorescent microscope with ApoTome.2.

### Homology-directed repair (HDR) genome editing

The donor ssAAV vector used as a template for HDR was constructed by cloning 1965bp GNA13 genomic DNA sequence flanking the 5’ and 3’ end of the target mutation into the AAV transfer plasmid between AAV ITR sequences. Control vector was generated with a synonymous substitution preserve the same codon QQQ. During genomic amplification, the AAA mutation was engineered: QQQ AAA (CAGCAACAG/GCTGCTGCT) within exon 4, and control synonymous substitution QQQ (CAGCAACAG/CAGCAGCAG) was generated in parallel. Additionally, in order to prevent endonuclease mediated re-cutting, the following PAM mutations were engineered, resulting in synonymous amino acid substitutions: C/T (YY) in exon 4. Primers contained homology arms to the AAV transfer vector allowing In-Fusion assembly into NheI/EoRI digested AAV plasmid. After confirmation of the insert sequence integrity, constructs were used with helper vectors from Cell Biolabs (VKP-400-DJ) for AAV virus production in Hek293t cells. The viruses were purified using AAVpro® Purification Kit Midi.

The guide sequences targeting GNA13 for CRISPR/Cas9 genome editing were predicted using the Feng Zhang lab website (Ref genome GRCh38, CRISPRko, SpyoCas9, Hsu design) and were ordered from IDT as sgRNA. The sgRNA: 5’-TGATAGCAGTGGTGAAGTGG-3’ was used in this study. For CRISPR/Cas9 genome editing, 73pmol of sgRNA was complexed with 61pmol recombinant Alt-RspCas9 protein in 8.31μL of PBS for 15 min and immediately used for nucleofection of 8×10^5^ primary keratinocytes with the Amaxa nucleofection apparatus (Lonza) using program T-018. After recovery, cells were mixed with AAV virus containing either control wild type QQQ or mutant donor QQQ AAA template at MOI 8×10^6^, transferred into 10cm dishes and cultured for minimal 72 hrs. Genomic DNAs were isolated and evaluated for the editing efficiency using PCR amplification and sanger sequencing of the bulk cell population.

For genomic DNA amplification, the following primers were used:

F: 5’-CCCATTGCTTTTAATAGAGGAGCA-3’

R: 5’-GGACTGGACAGGACAGCAAA-3’

### Quantitative RT-PCR analysis and primers

Total RNA was extracted using RNeasy Plus and further subjected to reverse transcription using the iScript cDNA Synthesis Kit. qRT-PCR analysis was performed using the LightCycler 480 II System (Roche) with the SYBR Green Master Mix. All Ct values are normalized to the levels of 60S ribosomal protein L32. The qRT-PCR primers are listed below.

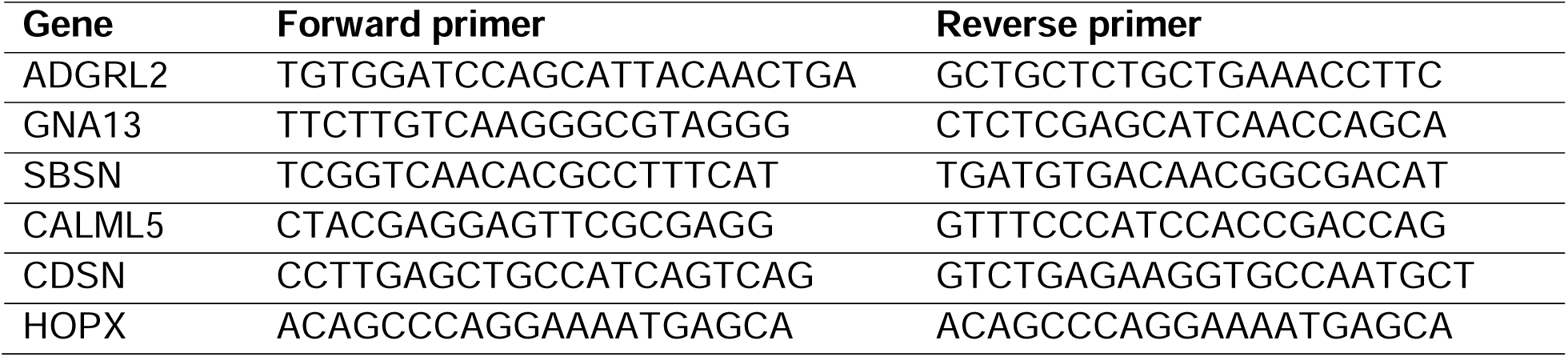
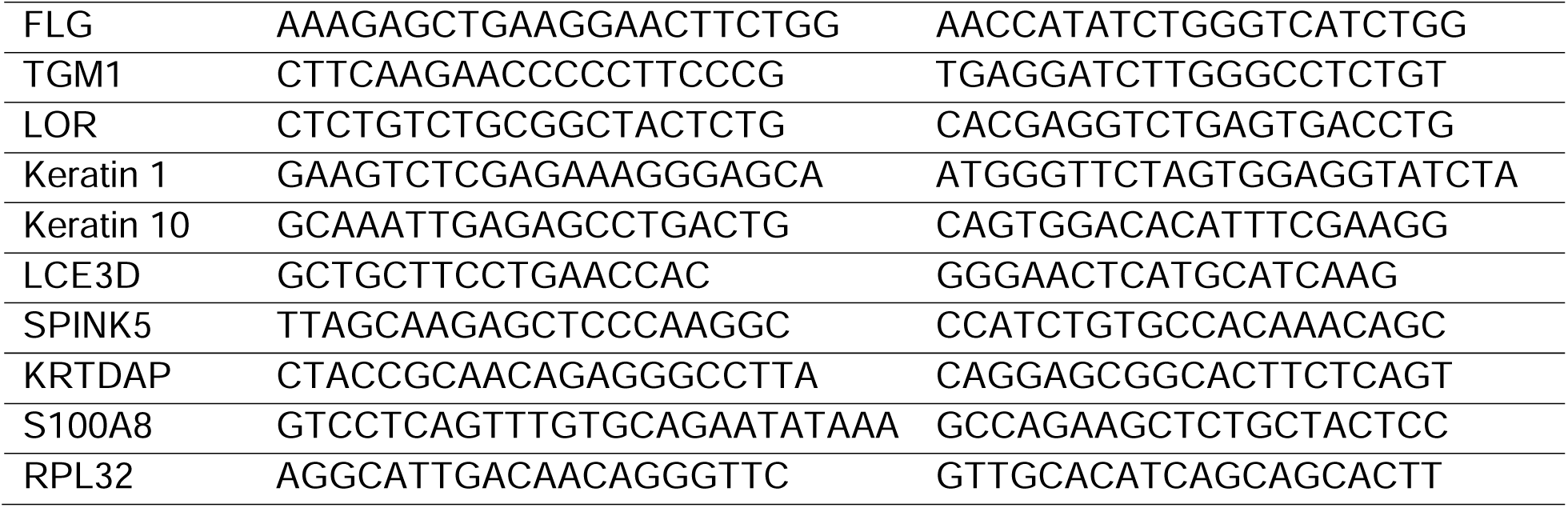

### Statistical analysis

Statistical tests were performed using GraphPad version 10.2.3. Sample sizes, statistical analyses methods, significance values are reported in the figure panel or figure legends. For statistical analyses, P > 0.05 were considered not significant (ns), and asterisks denote the following: * 0.01 < p ≤ 0.05; ** 0.001 < p ≤ 0.01; *** p ≤ 0.001. Error bars represent standard error mean (SEM) unless otherwise indicated.

### Reagent sources

| REAGENT | SOURCE | IDENTIFIER |
| --- | --- | --- |
| **Antibodies** |  |  |
| anti-ADGRL2 | Thermo Fisher Scientific | PA565359 |
| anti-GNA13 | proteintech | 67188-1-Ig |
| anti-Keratin 10 | Thermo Fisher Scientific | MA1-06319 |
| anti-HA | CST | 3724 |
| anti-Transglutaminase-1 | proteintech | 12912-3-AP |
| anti-Ki67 | Abcam | 15580 |
| anti-beta-actin | Millipore Sigma | A2228 |
| Anti Cytokeratin 10 Antibody (KRT10/844), Alexa Fluor™ 647 | Novus Biologicals | NBP2-47825AF647 |
| **Cell Lines** |  |  |
| Human primary keratinocytes | Stanford University School of Medicine | Genomic data sharing (GDS) |
| LentiX 293T | Takara Bio | 632180 |
| GKO HEK293 | Kind gift from Asuka Inoue, Tokyo University | Alvarez-Curto et al.,(19) J Biol Chem. Grundmann et al.,(18) Nat Commun. |
| **Reagents and Materials** |  |  |
| DMEM | Gibco | 11995-065 |
| Keratinocyte-SFM | Gibco | 17005-142 |
| Medium 154 | Gibco | M-154-500 |
| HKGS | Thermo Fisher Scientific | S0015 |
| rEGF | Gibco | 10450-013 |
| BPE | Gibco | 13028-014 |
| Antibiotic-Antimycotic | Thermo Fisher Scientific | 15240-062 |
| Penicillin-Streptomycin | Thermo Fisher Scientific | 15140122 |
| Dermis | New York Firefighter Skin Bank | N/A |
| Lipofectamine 3000 | Invitrogen | L3000015 |
| Lipofectamine RNAiMAX | Invitrogen | 13778150 |
| Lenti-X concentrator | Takara Bio | 631231 |
| RNeasy Plus Kit | QIAGEN | 74136 |
| iScript cDNA Synthesis Kit | Bio-Rad | 1708891 |
| SYBR Green Master Mix | Thermo Fisher Scientific | K0223 |
| QuantSeq 3' mRNA-seq Library Prep Kit FWD for Illumina | Lexogen | 015.96 |
| LDS sample loading buffer | Invitrogen | NP0008 |
| Opti-MEM | Thermo Fisher Scientific | 31985062 |
| Dispase | Corning | 354235 |
| Antibiotic-Antimycotic | Thermo Fisher Scientific | 15240-062 |
| Stellar Competent Cells | Takara Bio | 636763 |
| Thrombin | Sigma | T4846 |
| Enterokinase, light chain | NEB | P8070S |
| SPRIselect beads | Beckman Coulter | B23317 |
| RIPA Lysis Buffer | Thermo Fisher Scientific | 89900 |
| cOmplete™, Mini Protease Inhibitor Cocktail | Roche | 11836153001 |
| Fetal Bovine Serum | Gibco | 16000044 |
| Gibson Assembly Master Mix | NEB | E2611S |
| 4% E-Gel™ EX Agarose Gel | Thermo Fisher Scientific | G401004 |
| KAPA Library Quant Kit | Kapa Biosystems | KK4854 |
| AAVpro® Purification Kit Midi | Takara Bio | 6675 |
| AAV-DJ helper free packaging system | Cell Biolabs | VPK-400-DJ |
| MiSeq® Reagent Kit v3 150-cycle kit | Illumina | MS-102-3001 |
| cell strainer snap cap | VWR | 352235 |
| RNAseA | Qiagen | 19101 |
| Passive Lysis Buffer | Promega | E1910 |
| Proteinase K | Thermo Fisher Scientific | AM2546 |
| PrimeStar | Takara | R045A |
| Dual-Luciferase Reporter Assay System | Promega | E1910 |
| Non-essential Amino Acid Solution | Sigma | M7145 |
| Versene | Gibco | 15040066 |
| Hanks' balanced Salt Solution | Gibco | 14175095 |
| Coelenterazine-400a | NanoLight Technologies | 340 |
| Thrombin | Sigma | T4648 |
| Enterokinase | New England Biolabs | P8070S |
| BCA assay | Thermo Fisher Scientific | 23225 |
| 4% paraformaldehyde | Alfa Aesar | 43368 |
| DAPI | Sigma | DUO82040-5ML |
| Alt-RspCas9 protein | IDT | 10008100 |
| **Recombinant DNA** |  |  |
| pCMV-dR8.91 | Addgene | Romain Zufferey et al.,(60)<br>Nat. Biotechnol. |
| pMD2.G | Addgene | Romain Zufferey et al.,(60)<br>Nat. Biotechnol. |
| TRUPATH kit | Addgene | 1000000163 |
| SRF-RE reporter plasmid | Promega | E1350 |
| pRL-Renilla plasmid | Promega | E2261 |
| pLEX CMV Cas9 blast | This study | N/A |
| F+E 10x cs1 sgRNA mCherry | This study | N/A |
| Plex Flag-Keratin14 | This study | N/A |
| Plex HA-Keratin14 | This study | N/A |
| Plex full-length ADGRL2-FHH | This study | N/A |
| Plex ADGRL2-CTF-FHH | This study | N/A |
| Plex PAR1-ADGRL2-CTF-FHH | This study | N/A |
| Plex SNAP-Flag-ADGRL2-CTF-FHH | This study | N/A |
| pEB PAR-Lphn3tv7 | Gift from Richard Sando's lab | Duy Lan Huong Bui et al.,(20)<br>Cell Rep. |
| Plex ADGRL2-CTF-F943A-FHH | This study | N/A |
| Plex ADGRL2-CTF-V943A/F943A-FHH | This study | N/A |
| Plex full-length rat ADGRL2-FHH | This study | N/A |
| Plex full-length rat ADGRL2 F943A-FHH | This study | N/A |
| Plex Gα13-BRET | This study | N/A |
| Plex Gα13-AAA-BRET | This study | N/A |

## Data availability

All processed and raw sequencing data have been deposited in Gene Expression Omnibus (GEO), under the accession code: GSE281860. The Cryo-EM density maps and corresponding coordinates have been deposited in the Electron Microscopy Data Bank (EMDB) and the Protein Data Bank (PDB), respectively, under the following accession codes: EMD-47522 and 9E51.

## Author Contributions

X.Y. and P.A.K. conceived the project. F.H. and B.S. performed Cryo-EM, data collection and analysis, D.F.P., R.M.M., L.D. performed and supported for data analysis. K.G., D.L.H.B. performed TRUPATH assay. D.R., A.H. assisted with dual screening process. Z.S. assisted with HDR experiment. S.M. performed the luciferease assay. V.L.-P assisted the organoid study. K.L., T.S., J.Y.Q assisted with cell culture and cloning experiments. Y.X., F.H., G.S., and P.A.K. guided overall experiments and data analysis. X.Y. and P.A.K. wrote the manuscript with input from all authors.

## Acknowledgments

We thank T.C. Südhof, B.K. Kobilka, A.E. Oro, and H.Y. Chang for expert project advice. This work was supported by a USVA Merit Review grant BX001409 to P.A.K. and by NIAMS/NIH grants AR049737 to P.A.K. We thank Duy Thanh Nguyen, Ron Shanderson, Weili Miao, Ian Ferguson, Mårten Winge, Jordan Meyers, Jerry Qu, Joy Ji for the help and consultation during the project.

---

**Fig. S1.**
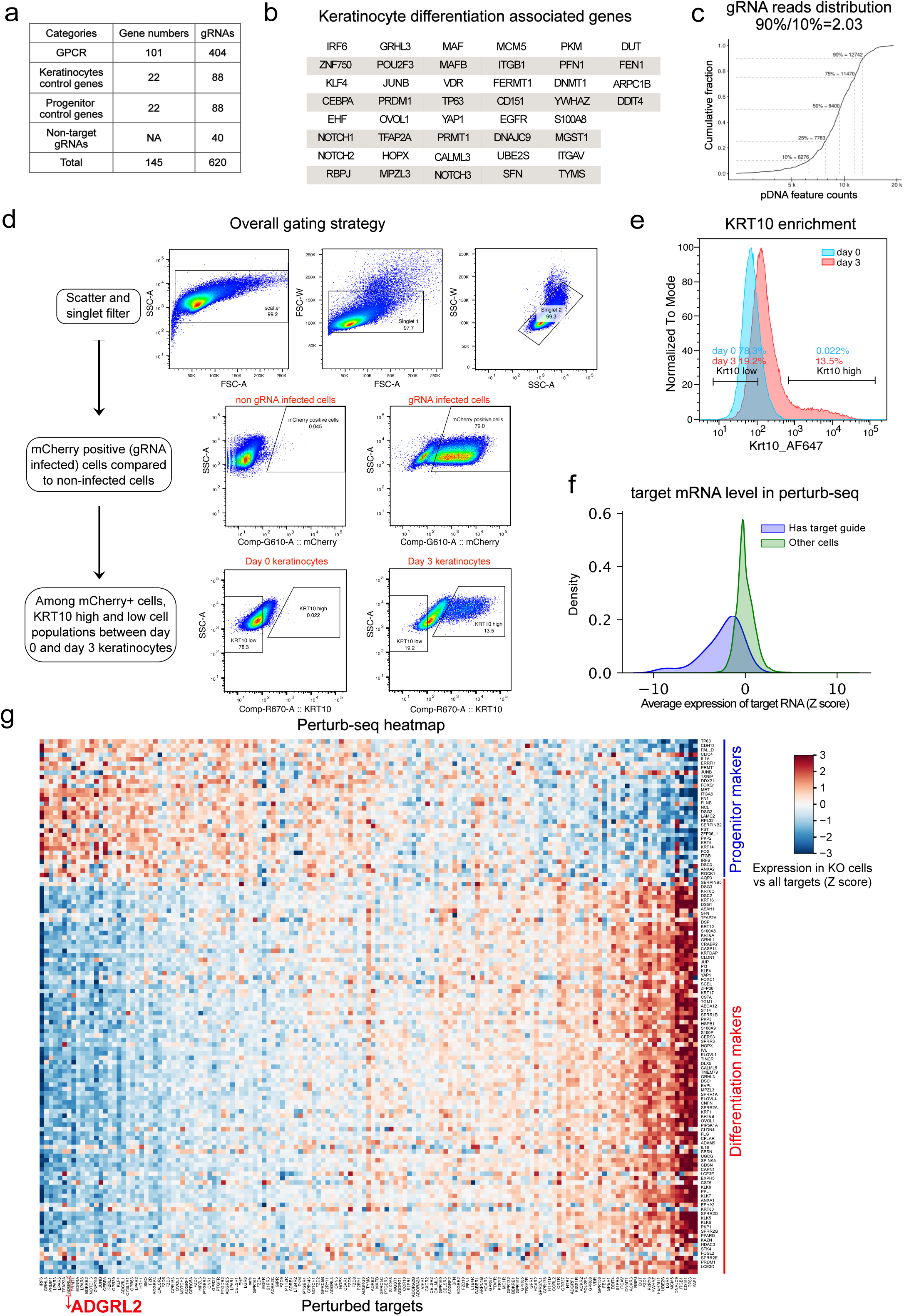
CRISPR-Flow and Perturb-seq GPCR knockout screens of epidermal GPCRs in differentiating keratinocytes. **a**, Numbers of gene targets and sgRNAs in the CRISPR plasmid library. **b**, Selected keratinocyte progenitor and differentiation genes. **c**, sgRNA count distribution in plasmid library. **d**, CRISPR-Flow gating strategy. **e**, Quantitative flow cytometric analysis of KRT10 expression changes from progenitors to differentiated cells. **f**, Perturb-seq mRNA overall knockout efficiency of target sgRNAs populations compared to non-target sgRNA populations. **g**, Heatmap visualization of Perturb-seq results; columns correspond to mRNAs while rows indicate the effect of individual gene perturbation on progenitor and differentiation genes.

**Fig. S2.**
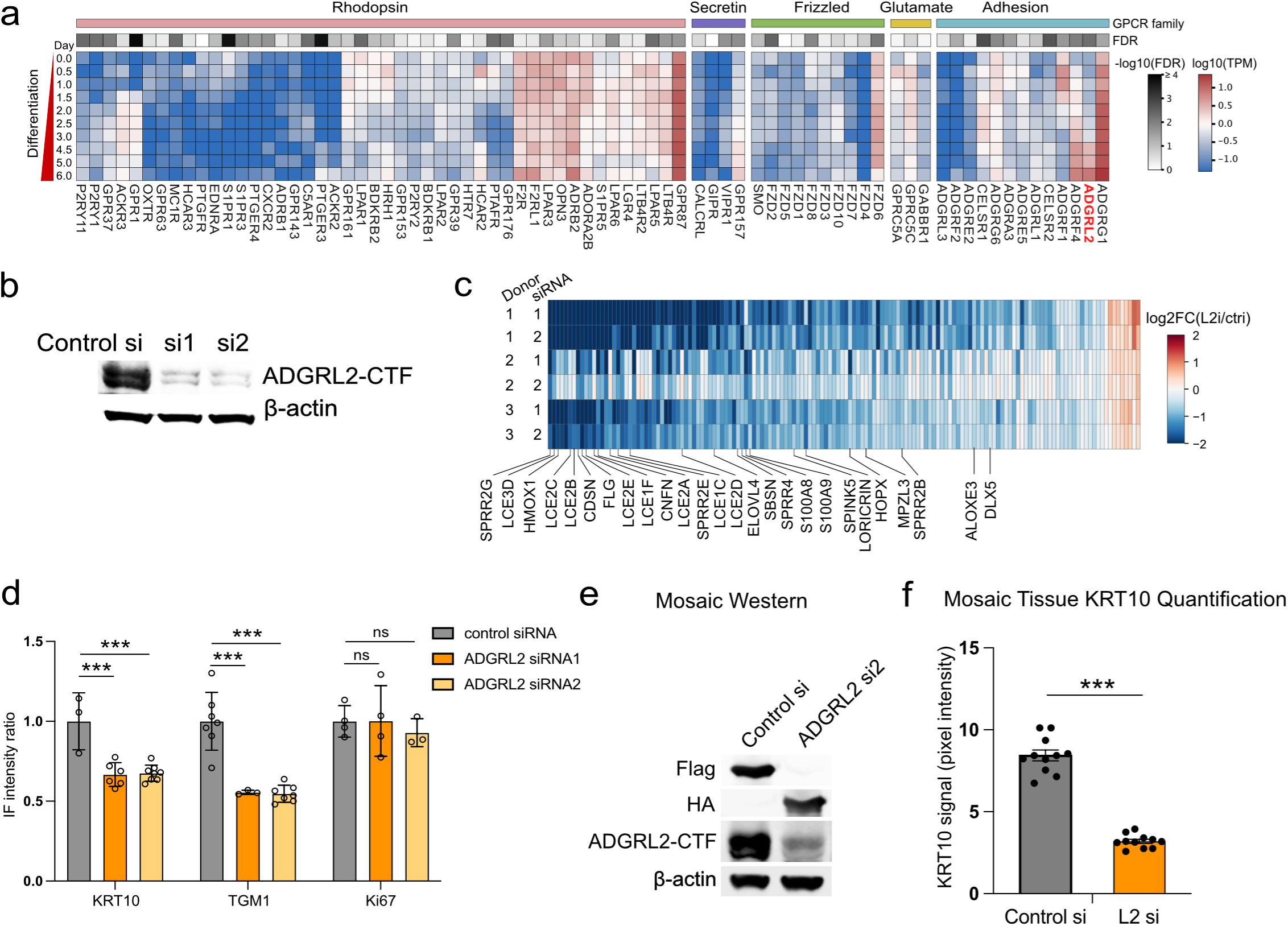
ADGRL2 affects keratinocyte differentiation in 2D and organoid models. **a**, Temporal heatmap illustrating GPCR mRNA expression during across days 0 to 6 of calcium-induced differentiation of primary human keratinocytes in vitro. **b**, ADGRL2 knockdown evaluation by Western blot. **c**, Heatmap of significant altered genes, FDR<0.05 with fold change >2 or <0.5, upon ADGRL2 knockdown. **d**, Quantification of KRT10, TGM1, Ki67 immunofluorescent signal in ADGRL2 knockdown organoid models, unpaired t Test. **e**, Western blot of the mosaic samples. **f**, Quantification of KRT10 signal in mosaic tissue control and L2 siRNAs. In **f**, paired t Test, n = 11.

**Fig. S3.**
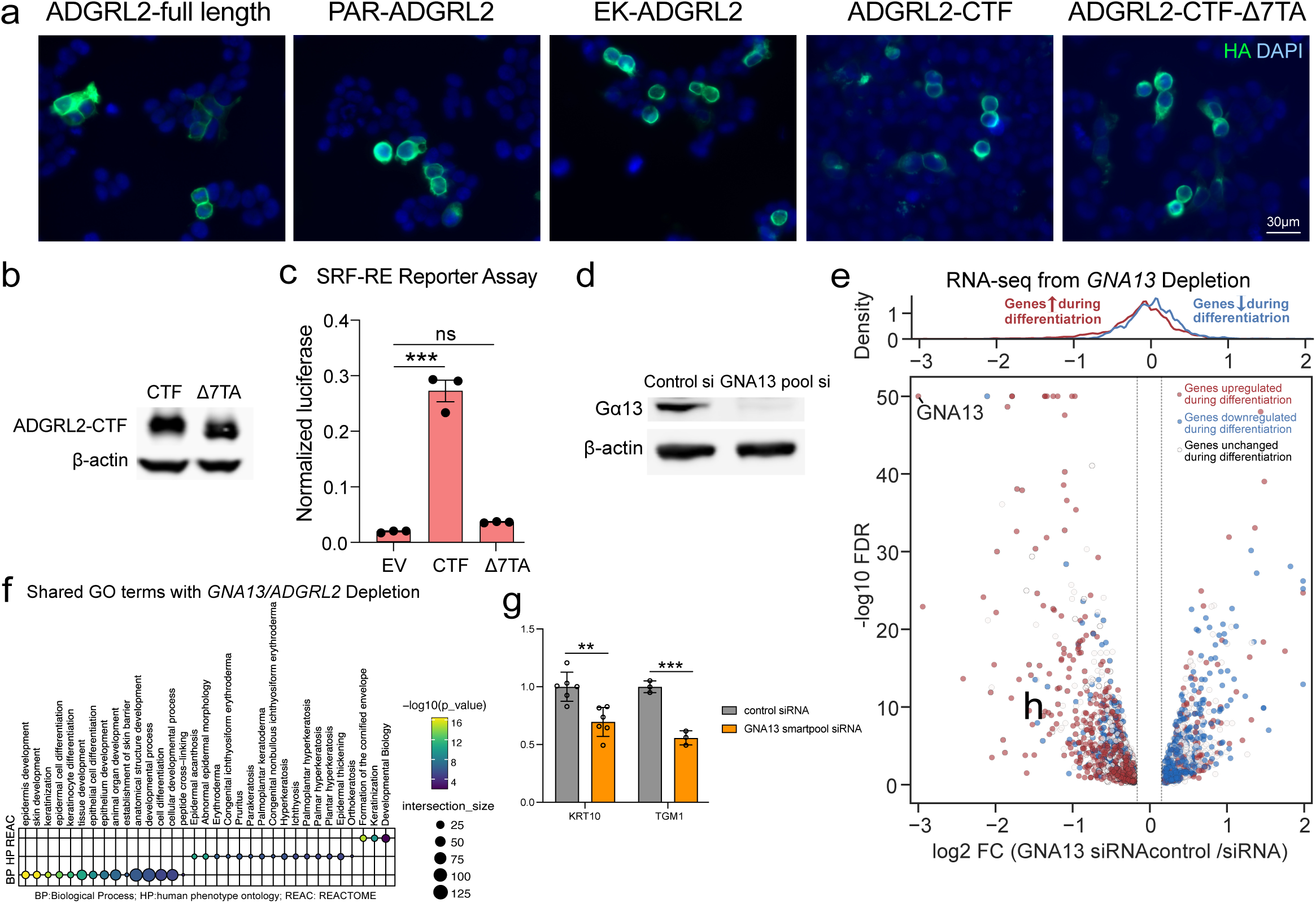
ADGRL2 activates Gα13 subtype to enable epidermal differentiation. **a**, Cellular localization of HA tagged ADGRL2 full-length, ADGRL2-CTF, PAR1-ADGRL2, EK-ADGRL2, ADGRL2-CTF-Δ7TA proteins in HEK293T cells evaluated by immunostaining. **b**, Western blot of ADGRL2-CTF and ADGRL2-CTF-Δ7TA protein expression. **c**, SRE-RF luciferase reporter measurements of ADGRL2-CTF and ADGRL2 TA peptide-deficient mutants. **d**, Gα proteins Perturb-seq showing enrichment of knock-out cells along the pseudotime trajectory (Mann-Whitney log10 P) compared to cells with safe target guides. A positive enrichment in low pseudotime cells indicates nominates a gene as necessary for differentiation. **e**, GNA13 knockdown evaluation by Western blot. **f**, RNA-Seq analysis of GNA13 knockdown samples sourced from two independent skin donors, showing all genes with log2 fold change >0.2 or <-0.2, red/blue genes are the ones upregulated/downregulated during differentiation, white genes are unchanged ones. **g**, Heatmap of significant altered genes, FDR<0.05 with fold change >2 or <0.5, upon GNA13 knockdown. **h**, Gene Ontology analysis of all significantly altered mRNAs (FDR<0.05) by GNA13 knockdown, based on RNA-seq data. **i**, Correlation between GNA13 and ADGRL2 knockdown RNA-seq results. **j**, Quantification of KRT10, TGM1 immunofluorescent signal in *GNA13* depleted regenerated human skin organoid tissue, unpaired t Test.

**Fig. S4.**
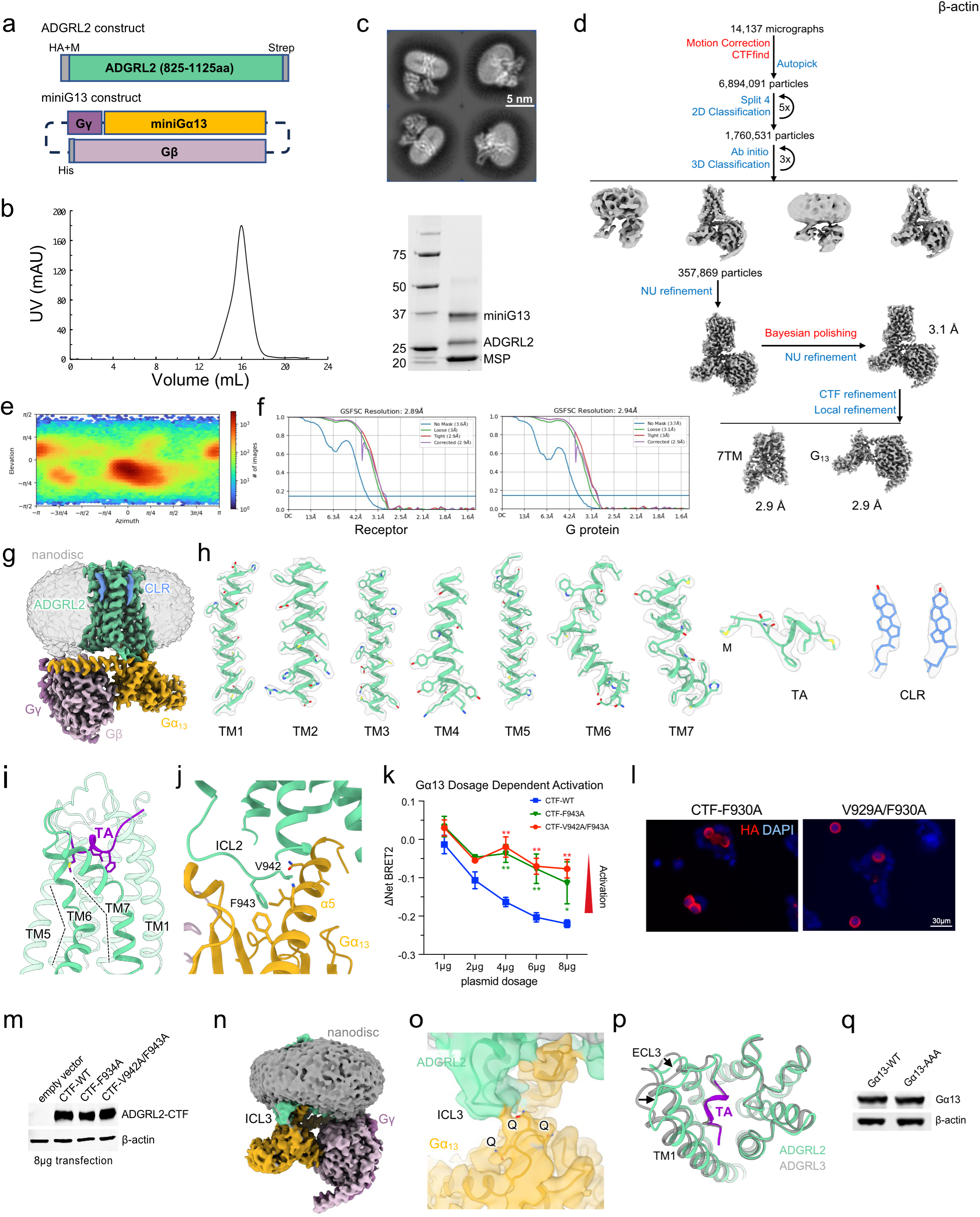
Cryo-EM analysis of the ADGRL2-Gα13 complex in lipid nanodiscs. **a**, The receptor and Gα13 constructs used in the study. Sequence corresponding to the CTF of ADGRL2 (TA and 7TM) was inserted after a hemagglutinin signal peptide (HA) and a methionine residue. The plasmid for expressing the miniGα13heterotrimer is the same as for the ADGRL3-Gα13 complex^37^. **b**, Size-exclusion chromatography (SEC) profile and SDS-PAGE of purified miniGα13-coupled ADGRL2. **c**, Representative 2D class averages of the ADGRL2-Gα13 complex. **d**, Cryo-EM data processing workflow for the ADGRL2-Gα13 complex in lipid nanodiscs. **e**, Angular distribution heat map of particles used for the global 3D reconstruction of the ADGRL3-Gα13 complex. **f**, Gold-standard Fourier shell correlation (FSC) curves of the locally refined receptor and miniGα13 reconstructions. **g**, Cryo-EM map of the ADGRL2-Gα13 complex in lipid nanodiscs showing the resolved density for two cholesterol molecules (colored in blue). The first methionine residue (M) used in the receptor construct was also resolved. **h**, Cryo-EM density and models are shown for TMs 1-7, the tethered agonist (TA) peptide and the CLR molecules of the ADGRL2-Gα13 complex. **i**, Binding mode of the TA peptide, and the bent TM6 and TM7 (highlighted by dashed lines) in the miniGα13-coupled ADGRL2. **j**, Interactions between ICL2 and the miniGα13 α5 helix of ADGRL2. Key residues are shown as sticks. **k**, Comparison of the basal Gα13 activity of ADGRL2-CTF WT and ICL2 mutants (F943A and V942/F943A) by titrating different amounts of transfected plasmids by BRET2 assay. **l**, Cellular localization of HA tagged ADGRL2-CTF-F943A and V942/F943A mutants in HEK293T cells evaluated by immunostaining. **m**, Western blot of empty vector, ADGRL2-CTF, ADGRL2-CTF-F943A and ADGRL2-CTF-V942/F943A in HEK293T cells. **n**, Unsharpened Cryo-EM map of the ADGRL2-Gα13 complex showing that ICL3 of the receptor protrudes from the lipid bilayer to interact with miniGα13. **o**, The model and EM density for the interface between ICL3 of the receptor and the QQQ patch of Gα13. **p**, Structural comparison of ADGRL2 and ADGRL3 when coupled with miniGα13, highlighting extracellular regions. **q**, Western blot of Gα13 wild-type and AAA mutant in HEK293T cells.

## Reference

1. V. Ratushny, M. D. Gober, R. Hick, T. W. Ridky, J. T. Seykora, From keratinocyte to cancer: the pathogenesis and modeling of cutaneous squamous cell carcinoma. J Clin Invest 122, 464–472 (2012).

2. X. Zhang, M. Yin, L. J. Zhang, Keratin 6, 16 and 17-Critical Barrier Alarmin Molecules in Skin Wounds and Psoriasis. Cells 8 (2019).

3. I. Pastar et al., Epithelialization in Wound Healing: A Comprehensive Review. Adv Wound Care (New Rochelle) 3, 445–464 (2014).

4. K. Alhosaini, A. Azhar, A. Alonazi, F. Al-Zoghaibi, GPCRs: The most promiscuous druggable receptor of the mankind. Saudi Pharm J 29, 539–551 (2021).

5. W. I. Weis, B. K. Kobilka, The Molecular Basis of G Protein–Coupled Receptor Activation. Annual Review of Biochemistry 87, 897–919 (2018).

6. R. Gutzmer, J. A. Solomon, Hedgehog Pathway Inhibition for the Treatment of Basal Cell Carcinoma. Target Oncol 14, 253–267 (2019).

7. A. Sekulic et al., Efficacy and safety of vismodegib in advanced basal-cell carcinoma. N Engl J Med 366, 2171–2179 (2012).

8. A. Ghahramani, G. Donati, N. M. Luscombe, F. M. Watt, Epidermal Wnt signalling regulates transcriptome heterogeneity and proliferative fate in neighbouring cells. Genome Biol 19, 3 (2018).

9. A. Sumitomo et al., LPA Induces Keratinocyte Differentiation and Promotes Skin Barrier Function through the LPAR1/LPAR5-RHO-ROCK-SRF Axis. J Invest Dermatol 139, 1010–1022 (2019).

10. S. Chen et al., Genome-wide CRISPR screen in a mouse model of tumor growth and metastasis. Cell 160, 1246–1260 (2015).

11. K. J. Condon et al., Genome-wide CRISPR screens reveal multitiered mechanisms through which mTORC1 senses mitochondrial dysfunction. Proc Natl Acad Sci U S A 118 (2021).

12. M. Fomicheva, I. G. Macara, Genome-wide CRISPR screen identifies noncanonical NF-kappaB signaling as a regulator of density-dependent proliferation. Elife 9 (2020).

13. B. Adamson et al., A Multiplexed Single-Cell CRISPR Screening Platform Enables Systematic Dissection of the Unfolded Protein Response. Cell 167, 1867–1882 e1821 (2016).

14. A. Dixit et al., Perturb-Seq: Dissecting Molecular Circuits with Scalable Single-Cell RNA Profiling of Pooled Genetic Screens. Cell 167, 1853–1866 e1817 (2016).

15. D. S. Kim et al., The dynamic, combinatorial cis-regulatory lexicon of epidermal differentiation. Nat Genet 53, 1564–1576 (2021).

16. K. Cockburn et al., Gradual differentiation uncoupled from cell cycle exit generates heterogeneity in the epidermal stem cell layer. Nat Cell Biol 24, 1692–1700 (2022).

17. R. H. J. Olsen et al., TRUPATH, an open-source biosensor platform for interrogating the GPCR transducerome. Nat Chem Biol 16, 841–849 (2020).

18. M. Grundmann et al., Lack of beta-arrestin signaling in the absence of active G proteins. Nat Commun 9, 341 (2018).

19. E. Alvarez-Curto et al., Targeted Elimination of G Proteins and Arrestins Defines Their Specific Contributions to Both Intensity and Duration of G Protein-coupled Receptor Signaling. J Biol Chem 291, 27147–27159 (2016).

20. D. L. H. Bui et al., The adhesion GPCRs CELSR1-3 and LPHN3 engage G proteins via distinct activation mechanisms. Cell Rep 42, 112552 (2023).

21. P. Xiao et al., Tethered peptide activation mechanism of the adhesion GPCRs ADGRG2 and ADGRG4. Nature 604, 771–778 (2022).

22. S. Mathiasen et al., G12/13 is activated by acute tethered agonist exposure in the adhesion GPCR ADGRL3. Nature Chemical Biology 16, 1343–1350 (2020).

23. N. A. Perry-Hauser, M. W. VanDyck, K. H. Lee, L. Shi, J. A. Javitch, Disentangling autoproteolytic cleavage from tethered agonist-dependent activation of the adhesion receptor ADGRL3. J Biol Chem 298, 102594 (2022).

24. D. T. Pederick et al., Context-dependent requirement of G protein coupling for Latrophilin-2 in target selection of hippocampal axons. Elife 12 (2023).

25. H. M. Stoveken, A. G. Hajduczok, L. Xu, G. G. Tall, Adhesion G protein-coupled receptors are activated by exposure of a cryptic tethered agonist. Proc Natl Acad Sci U S A 112, 6194–6199 (2015).

26. X. Barros-Alvarez et al., The tethered peptide activation mechanism of adhesion GPCRs. Nature 604, 757–762 (2022).

27. Y. Q. Ping et al., Structural basis for the tethered peptide activation of adhesion GPCRs. Nature 604, 763–770 (2022).

28. X. Qu et al., Structural basis of tethered agonism of the adhesion GPCRs ADGRD1 and ADGRF1. Nature 604, 779–785 (2022).

29. Y. Qian et al., Structural insights into adhesion GPCR ADGRL3 activation and G(q), G(s), G(i), and G(12) coupling. Mol Cell 82, 4340–4352 e4346 (2022).

30. D. T. D. Jones et al., Tethered agonist activated ADGRF1 structure and signalling analysis reveal basis for G protein coupling. Nat Commun 14, 2490 (2023).

31. M. M. Papasergi-Scott et al., Time-resolved cryo-EM of G-protein activation by a GPCR. Nature 629, 1182–1191 (2024).

32. M. P. Pedro et al., GPCR Screening Reveals that the Metabolite Receptor HCAR3 Regulates Epithelial Proliferation, Migration, and Cellular Respiration. J Invest Dermatol 144, 1311–1321 e1317 (2024).

33. W. C. Probst, L. A. Snyder, D. I. Schuster, J. Brosius, S. C. Sealfon, Sequence alignment of the G-protein coupled receptor superfamily. DNA Cell Biol 11, 1–20 (1992).

34. V. Katritch, V. Cherezov, R. C. Stevens, Diversity and modularity of G protein-coupled receptor structures. Trends Pharmacol Sci 33, 17–27 (2012).

35. A. J. Venkatakrishnan et al., Structured and disordered facets of the GPCR fold. Curr Opin Struct Biol 27, 129–137 (2014).

36. I. G. Denisov, S. G. Sligar, Nanodiscs for structural and functional studies of membrane proteins. Nat Struct Mol Biol 23, 481–486 (2016).

37. F. He et al., Allosteric modulation and G-protein selectivity of the Ca(2+)-sensing receptor. Nature 626, 1141–1148 (2024).

38. F. Sadler et al., Autoregulation of GPCR signalling through the third intracellular loop. Nature 615, 734–741 (2023).

39. R. H. Purcell, R. A. Hall, Adhesion G Protein-Coupled Receptors as Drug Targets. Annu Rev Pharmacol Toxicol 58, 429–449 (2018).

40. R. Sando, T. C. Sudhof, Latrophilin GPCR signaling mediates synapse formation. Elife 10 (2021).

41. R. Sando, X. Jiang, T. C. Sudhof, Latrophilin GPCRs direct synapse specificity by coincident binding of FLRTs and teneurins. Science 363 (2019).

42. A. Krishnan, S. Nijmeijer, C. de Graaf, H. B. Schioth, Classification, Nomenclature, and Structural Aspects of Adhesion GPCRs. Handb Exp Pharmacol 234, 15–41 (2016).

43. S. L. Regan, M. T. Williams, C. V. Vorhees, Latrophilin-3 disruption: Effects on brain and behavior. Neurosci Biobehav Rev 127, 619–629 (2021).

44. M. L. Herman et al., Transglutaminase-1 gene mutations in autosomal recessive congenital ichthyosis: summary of mutations (including 23 novel) and modeling of TGase-1. Hum Mutat 30, 537–547 (2009).

45. J. van der Velden, M. van Geel, J. J. Engelhart, M. F. Jonkman, P. M. Steijlen, Mutations in the CDSN gene cause peeling skin disease and hypotrichosis simplex of the scalp. J Dermatol 47, 3–7 (2020).

46. E. Pohler et al., Novel autosomal dominant mutation in loricrin presenting as prominent ichthyosis. Br J Dermatol 173, 1291–1294 (2015).

47. L. Guerra et al., Ichthyosis Linearis Circumflexa as the Only Clinical Manifestation of Netherton Syndrome. Acta Derm Venereol 95, 720–724 (2015).

48. J. P. Thyssen, E. Godoy-Gijon, P. M. Elias, Ichthyosis vulgaris: the filaggrin mutation disease. Br J Dermatol 168, 1155–1166 (2013).

49. L. Kolberg, U. Raudvere, I. Kuzmin, J. Vilo, H. Peterson, gprofiler2 -- an R package for gene list functional enrichment analysis and namespace conversion toolset g:Profiler. F1000Res 9 (2020).

50. A. Peisley, G. Skiniotis, 2D Projection Analysis of GPCR Complexes by Negative Stain Electron Microscopy. Methods Mol Biol 1335, 29–38 (2015).

51. J. Zivanov et al., New tools for automated high-resolution cryo-EM structure determination in RELION-3. Elife 7 (2018).

52. S. Q. Zheng et al., MotionCor2: anisotropic correction of beam-induced motion for improved cryo-electron microscopy. Nat Methods 14, 331–332 (2017).

53. A. Rohou, N. Grigorieff, CTFFIND4: Fast and accurate defocus estimation from electron micrographs. J Struct Biol 192, 216–221 (2015).

54. A. Punjani, J. L. Rubinstein, D. J. Fleet, M. A. Brubaker, cryoSPARC: algorithms for rapid unsupervised cryo-EM structure determination. Nat Methods 14, 290–296 (2017).

55. E. F. Pettersen et al., UCSF Chimera--a visualization system for exploratory research and analysis. J Comput Chem 25, 1605–1612 (2004).

56. P. Emsley, B. Lohkamp, W. G. Scott, K. Cowtan, Features and development of Coot. Acta Crystallogr D Biol Crystallogr 66, 486–501 (2010).

57. D. Liebschner et al., Macromolecular structure determination using X-rays, neutrons and electrons: recent developments in Phenix. Acta Crystallogr D Struct Biol 75, 861–877 (2019).

58. V. B. Chen et al., MolProbity: all-atom structure validation for macromolecular crystallography. Acta Crystallogr D Biol Crystallogr 66, 12–21 (2010).

59. E. F. Pettersen et al., UCSF ChimeraX: Structure visualization for researchers, educators, and developers. Protein Sci 30, 70–82 (2021).

60. R. Zufferey, D. Nagy, R. J. Mandel, L. Naldini, D. Trono, Multiply attenuated lentiviral vector achieves efficient gene delivery in vivo. Nat Biotechnol 15, 871–875 (1997).

